# HyperVR–A hybrid prediction framework for virulence factors and antibiotic resistance genes in microbial data

**DOI:** 10.1101/2022.05.24.493218

**Authors:** Boya Ji, Wending Pi, Xianglilan Zhang, Shaoliang Peng

**Affiliations:** College of Computer Science and Electronic Engineering, Hunan University, Changsha 410006, People’s Republic of China; State Key Laboratory of Pathogen and Biosecurity, Beijing Institute of Microbiology and Epidemiology, Beijing 100071, People’s Republic of China

**Keywords:** virulence factors, antibiotic resistance genes, microbial data, machine learning, deep learning

## Abstract

Infectious diseases, particularly bacterial infections, are emerging at an unprecedented rate, posing a serious challenge to public health and the global economy. Different virulence factors (VFs) work in concert to enable pathogenic bacteria to successfully adhere, reproduce and cause damage to host cells, and antibiotic resistance genes (ARGs) allow pathogens to evade otherwise curable treatments. To understand the causal relationship between microbiome composition, function and disease, both VFs and ARGs in microbial data must be identified. Most existing computational models cannot simultaneously identify VFs or ARGs, hindering the related research. The best hit approaches are currently the main tools to identify VFs and ARGs concurrently; yet they usually have high false-negative rates and are very sensitive to the cut-off thresholds. In this work, we proposed a hybrid computational framework called HyperVR to predict VFs and ARGs at the same time. Specifically, HyperVR integrates key genetic features and then stacks classical ensemble learning methods and deep learning for training and prediction. HyperVR accurately predicts VFs, ARGs and negative genes (neither VFs nor ARGs) simultaneously, with both high precision (>0.91) and recall (>0.91) rates. Also, HyperVR keeps the flexibility to predict VFs or ARGs individually. Regarding novel VFs and ARGs, the VFs and ARGs in metagenomic data, and pseudo VFs and ARGs (gene fragments), HyperVR has shown good prediction, outperforming the current state-of-the-art predition tools and best hit approaches in terms of precision and recall. HyperVR is a powerful tool for predicting VFs and ARGs simultaneously by using only gene sequences and without strict cut-off thresholds, hence making prediction straightforward and accurate.

## Introduction

The microbiome is critical to the inner ecosystem of hosts, e.g., humans, higher animals and plants, as well as to maintain the external environment [1, 2, 3, 4]. In particular, pathogenic microorganisms cause diseases and even threaten the life of the host by carrying virulence factors (VFs) and/or antibiotic resistance genes (ARGs) [5, 6]. Accurate and timely identification of VFs and ARGs can effectively guide medical treatments, decrease host morbidity and mortality, and reduce the economic losses in husbandry, aquaculture, etc.

The VFs in pathogenic microorganisms can induce pathogenicity, with the ability to assist the microorganisms in colonising their hosts at the cellular level [7]. The success of pathogenic microorganisms leans on their power to utilize VFs to make infection, survive in the hostile host environment and cause the disease. The assorted expressions, organizations and combinations of VFs are answerable for different clinical symptoms of pathogenic infection [8]. VFs can be classified into various categories such as adhesion, colonization, exotoxin, endotoxin, iron transport, etc. The different VFs work in concert to enable the pathogenic microorganisms, such as bacteria and fungi, to successfully adhere, reproduce and cause damage to the host cells [9].

The ARGs are barriers to the treatment of pathogenic infections, exacerbating the pathogenic ability of microorganisms [10]. Antibiotics have proven effective in treating a variety of microbial infections, especially bacterial infections over the years [11]. However, the treatments for bacterial infections are increasingly limited, and the previous effective treatment options even do not exist for some patients. Particularly, an increasing number of antibiotic-resistant bacteria require complex infection treatments, and some infections, such as those caused by multi-drug resistant bacteria, are now unresponsive to conventional first-line treatments [12]. Some examples are *Vancomycin-Resistant Enterococci* (VRE), which is a global nosocomial pathogen with important clinical implications [13]; *Methicillin-Resistant Staphylococcus Aureus* (MRSA), which is the leading cause of hospital- and community-acquired infections, leading to severe mortality and morbidity [14]; and *Colistin-Carbapenem-Resistant Escherichia Coli* (CCREC), which develops resistance to the last-resort drug by acquiring ARGs *bla*_*NDM*–1_ and *mcr-1* [15].

To understand the causal relationship between microbiome composition, function, and disease, both VFs and ARGs must be identified. Despite the different evolutionary pathways, VFs and ARGs have common features that are necessary for pathogenic bacteria to adapt to and survive in a competitive microbial environment. Specifically, both VFs and ARGs are frequently transferred between bacteria through horizontal gene transfer (HGT) and both utilize similar systems (i.e., two-component systems, efflux pumps, cell wall alterations, and porins) to activate or repress the expression of various genes [16]. Pathogens can use VFs to cause diseases in their hosts, while they can colonise an environment with selective antibiotic pressure through the acquisition or presence of ARGs [17, 18, 19].

The best hit approaches are currently the primary means to simultaneously identify VFs and ARGs by comparing metagenomic gene sequences with existing online databases using programs such as BLAST [20], DIAMOND [21], or Bowtie [22]. With the advent of high-throughput sequencing technologies, powerful and sensitive metagenomic approaches are available for the analysis of small molecules, proteins, and nucleic acids (DNA and RNA) obtained from biological samples in a wide range of environmental compartments, such as the human microbiome, water, soil, compost, livestock manure, wastewater treatment plants, etc [23, 24]. The best hit approaches align predicted open reading frames from assembled contigs or raw reads to known gene databases obtained from the analysis and then uses an alignment length requirement or sequence similarity cut-offs to predict or assign classes of genes.

To our knowledge, little attention has been paid to the study of computational methods that can simultaneously identify/predict VFs and ARGs, and only a few tools that utilize the best hit approaches are available. In particular, VR-profile [25] enables the identification of VFs and/or ARGs and their transfer-associated gene clusters by performing homology searches on the genome sequences of pathogenic bacteria. It first collected a large number of known gene cluster loci of bacterial mobile genetic elements, including pathogenicity/antibiotic resistance islands, IS elements, class I integrons, prophages, and conjugative and integrative elements, to construct a MobilomeDB database. Then, HMMer or BLASTp was used to perform a fast homology search for the queried genomic sequences against MobilomeDB. Furthermore, the PathoFact [26] combined different existing modules and databases to build a pipeline for the identification of VFs, toxin genes and ARGs.

The best hit approaches typically have low false positive rates [24], i.e. few non-VFs and non-ARGs are predicted to be VFs and ARGs, but high false negative rates [27, 28], i.e. a large number of actual VFs and ARGs are predicted to be non-VFs and non-ARGs. In addition, these approaches are very sensitive to the cut-off threshold; in general, when the cut-off value is high, the predicted results have high precision but low recall, and when the cut-off value is low, the predicted results have high recall but low precision, making it impossible for novice bioinformaticians to decide on an appropriate cut-off threshold.

To address these limitations, we proposed a hybrid computational framework called HyperVR to predict VFs and ARGs at the same time. Specifically, HyperVR integrates multiple key genetic features, including bit score-based similarity feature, physicochemical property-based features, evolutionary information-based features, and one-hot encoding feature, and then stacks classical ensemble learning methods and deep learning for training and prediction (see Figure 1). To accurately evaluate the performance of HyperVR in predicting ARGs, VFs, and negative genes (neither VFs nor ARGs) simultaneously, a 5-fold cross-validation method was performed, resulting in a micro-average precision and recall of 91.94% with a standard deviation of 0.0065. To further evaluate the performance of HyperVR in predicting ARGs (HyperVR-ARGs) or VFs (HyperVR-VFs) individually, the same 5-fold cross-validation method was performed. Furthermore, we also evaluated the ability of HyperVR for novel VFs and ARGs, the VFs and ARGs in metagenomic data, and pseudo VFs and ARGs (gene frgments). Using the three simulated microbial datasets, we also performed a comparison of HyperVR to other baseline methods. As a result, HyperVR all showed better predictive capability, outperforming the current state-of-the-art predition tools and best hit approaches in terms of precision and recall. Finally, we selected three specific pathogen strain datasets to test the ability of HyperVR to simultaneously identify ARGs and VFs in real pathogenic bacteria.

**Figure 1.**
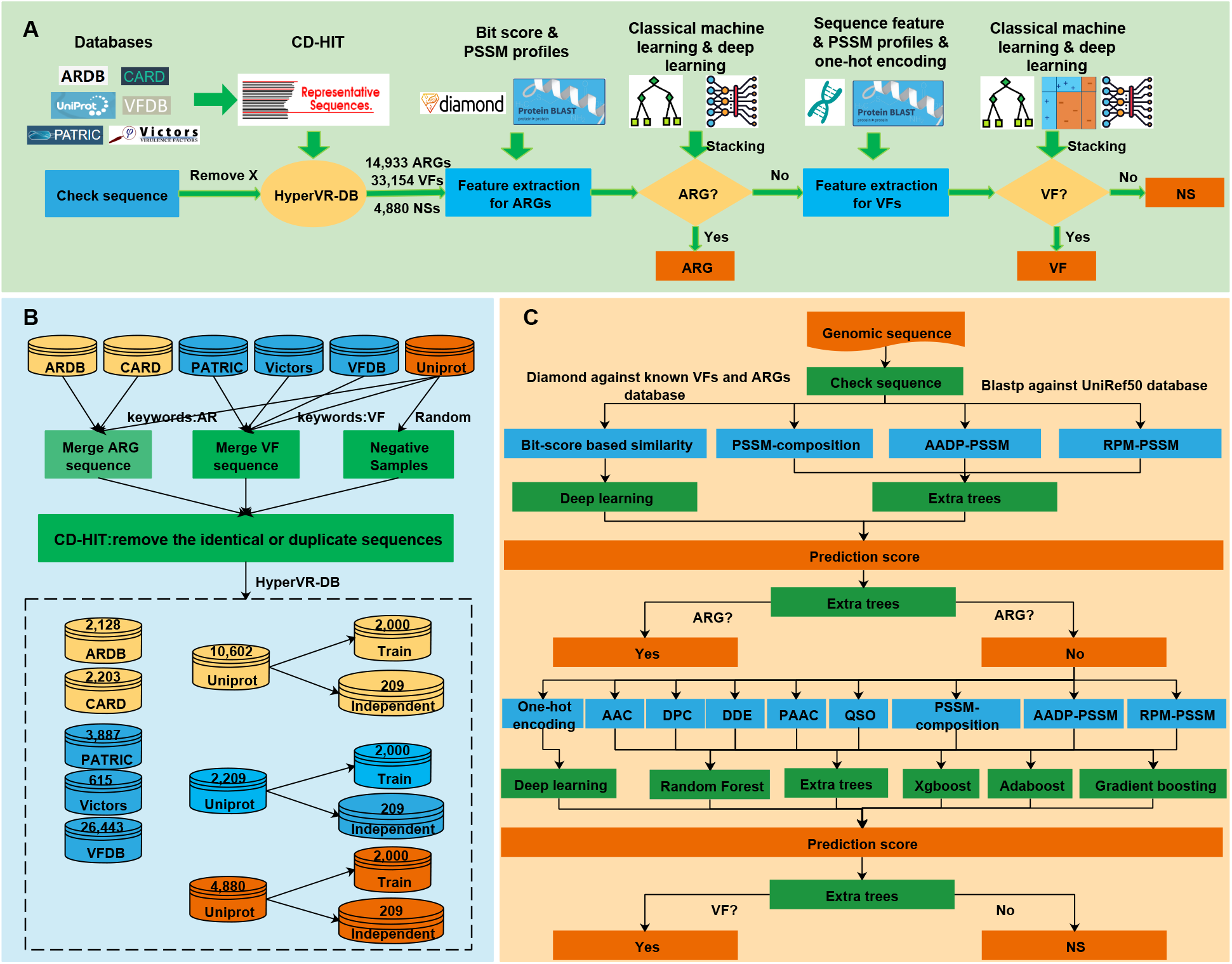
Schematic illustration of HyperVR for simultaneous prediction of ARGs and VFs. **A.** The overview of HyperVR. The first step was to collect and organize the dataset, including removing characters other than the 20 natural amino acid abbreviations from the sequences (remove X) and using CD-HIT to remove identical or duplicate sequences from the three types of samples to ensure the accuracy of the model prediction results. The second step is to predict whether the gene is an ARG, including feature extraction (generation of bit-score based similarity feature using the diamond program and evolutionary information-based features using the blast program) and stacking model training and prediction (classical ensemble learning and deep learning); if the judgment result is no, go to the third step; otherwise, output the prediction score of the gene belonging to ARGs. The third step is then to predict whether the gene is a VF, again including feature extraction (generation of sequence-based features, one-hot encoding feature and evolutionary information-based features using the blast program) and stacking model training and prediction (classical ensemble learning, boosting learning, and deep learning); if the judgment result is no, output the prediction score of the gene belonging to NSs; otherwise, output the prediction score of the gene belonging to VFs. **B.** The detailed procedures for data processing. Yellow represents the ARGs dataset, blue represents the VFs dataset, and orange represents the NSs dataset; 2000 ARGs, VFs and NSs in the UNIPROT database were used for model training and validation to address the data imbalance; 209 ARGs, VFs and NSs in the UNIPROT database were used for the independent test dataset. **C.** The detailed flowchart of HyperVR. The top half of the flowchart can be used individually to predict ARGs (HyperVR-ARG) and the bottom half of the flowchart can be used individually to predict VFs (HyperVR-VF).

## 1. Materials and Methods

### 1.1 Data collection and annotation

#### 1.1.1 Antibiotic Resistance Genes (ARGs)

The original ARGs in this work were firstly derived from DeepARG-DB [29], including three major databases: UNIPROT [30], CARD [31] and ARDB [32]. For the UNIPROT dataset, all genes containing the antibiotic resistance keyword (KW-0046) or a metadata description were selected and further annotated through a manual inspection and text mining, removing sequences annotated as conferring resistance by single nucleotide polymorphisms (SNPs). Secondly, all sequences from the three datasets (UNIPROT + CARD + ARDB) were clustered using CD-HIT [33], thereby removing all identical or duplicate sequences by setting the identity parameter to 100%. This step aims to make the following divided training dataset and known ARGs have no identical genes so that the final model training results are credible. Finally, 14,933 ARGs were retained for downstream analysis, including 10,602 UNIPROT, 2,203 CARD, and 2,128 ARDB genes.

#### 1.1.2 Virulence Factors (VFs)

The original VFs in this work were collected from four public databases: PATRIC [34], Victors [35], VFDB [8] and UNIPROT Swiss-Prot [36]. Specifically, we firstly downloaded 1,293, 4,964, and 28,913 VFs from PATRIC, Victors, and VFDB, respectively. For the UNIPROT Swiss-Prot dataset, 4,085 genes that contained the virulence factors keyword (KW-0843) were selected. Secondly, all sequences from the four datasets (PATRIC + Victors + VFDB + UNIPROT) were clustered using CD-HIT similar to the ARGs operation by setting the identity parameter to 100%. Finally, 33,154 VFs were retained for downstream analysis, including 3,887 PATRIC, 615 Victors, 26,443 VFDB, and 2,209 UNIPROT genes.

#### 1.1.3 Negative Samples (NSs)

For negative samples, i.e., genes that were neither ARGs nor VFs, we firstly random collected 20,000 genes from UNIPROT Swiss-Prot [36] (3.5% of the original UNIPROT Swiss-Prot genes). Secondly, all the ARGs and VFs mentioned earlier, and genes containing the virulence or antibiotic keywords (Supplementary File Table S1) were removed. Thirdly, the remaining genes and all the ARGs and VFs mentioned earlier were clustered using CD-HIT to ensure maximum cleanliness between the three classes of samples. Finally, 4,880 genes were obtained as negative samples.

#### 1.1.4 HyperVR-DB

The resulting database, HyperVR-DB, comprises 52967 genes, including 14,933 ARGs, 33,154 VFs, and 4,880 NSs. Next, 2000 ARGs, VFs and NSs in the UNIPROT database were used for model training and validation to address the data imbalance; 209 ARGs, VFs and NSs in the UNIPROT database were used for the independent test dataset; genes in the remaining database were used to represent known ARGs and VFs (see Figure 1.B).

### 1.2 Feature Extraction

To obtain a superior predictive power for ARGs and VFs, HyperVR considers a variety of gene-related features, including the following five categories: Bit score-based similarity feature, Sequence-based features, Physicochemical property-based features, Evolutionary information-based features, and One-hot encoding feature. Detailed feature descriptions are presented in the following subsections.

#### 1.2.1 Bit score-based similarity feature

The Bit score-based similarity feature [29] consists of the bit scores between full gene length sequences and known ARGs and VFs, which considers the similarity distribution of sequences in the ARGs and VFs databases, not just the best hits. The bit score is used as a similarity metric because it considers the identity extent between sequences and, unlike the e-value, it is independent of the size of the database [37]. In this work, we chose the DIAMOND program, which is faster than BLAST, to align the gene sequences in the training dataset with the remaining known 12724 (14933-2209) ARGs and 30945 (33154-2209) VFs used for comparison in HyperVR-DB under the more-sensitive parameter. It should be noted that the training dataset has been de-duplicated using CD-HIT program with the dataset used for comparison to avoid the possibility of label leakage (Refer to Section 1.1). Then, the bit scores are normalized to the [0, 1] interval to represent the similarity of the sequences in terms of distance. Finally, the Bit score-based similarity feature of each gene sequence in the training dataset is transformed into a fixed 12724+30945=43669-dimensional feature vector, where each dimension is the bit score output by DIAMOND program between full gene length sequences and each available ARGs and VFs in the comparison dataset.

#### 1.2.2 Sequence-based features

##### Amino Acid Composition (AAC)

The AAC feature [38] indicates the frequencies of 20 natural amino acids (i.e. “ACDEFGHIKLMNPQRSTVWY”) in a protein or peptide sequence and can be calculated as:

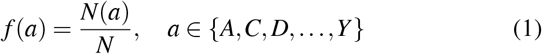

where *N*(*a*) denotes the number of a given amino acid *a, N* denotes the sequence length of the protein or peptide, and *f*(*a*) denotes the final generated 20-dimensional feature vector.

##### Di-Peptide Composition (DPC)

The DPC feature [39] indicates the frequencies of di-peptide in a protein or peptide sequence and can be calculated as:

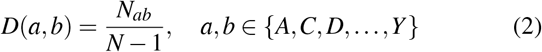

where *N_ab_* denotes the number of a given di-peptide *ab, N* denotes the sequence length of the protein or peptide, and *D*(*a, b*) denotes the final generated 20×20=400-dimensional feature vector.

##### Di-peptide Deviation from Expected Mean (DDE)

The DDE feature [39] is a combination of three features: theoretical mean (TM), di-peptide composition (DPC), and theoretical variance (TV). Specifically, the TM feature is calculated as follows:

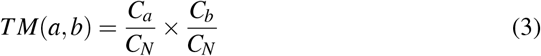

where *C_a_* and *C_b_* respectively denotes the codon numbers encoding amino acid *a* and *b*, and *C_N_* equals 61, denoting the total number of possible codons without including the three stop codons. The calculation of DPC feature refers to the previous description. The TV feature is calculated as follows:

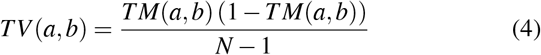

Finally, *DDE* (*a, b*) is calculated as follows:

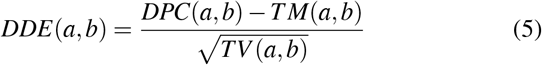

#### 1.2.3 Physicochemical property-based features

##### Pseudo-Amino Acid Composition (PAAC)

The PAAC feature [40, 41] contains two aspects. First, the original side chain masses, hydrophobicity, and hydrophilicity of the 20 natural amino acids are defined as *M^o^*(*i*), 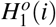, and 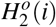 for *i* = 1,2,…,20, respectively. Second, they are normalized as follows:

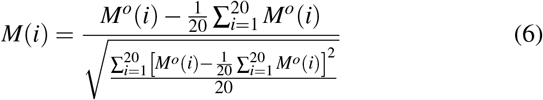

where 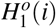 and 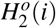 are normalized in the same way. Third, the correlation between *R_i_*, and *R_j_* that possess a set of *n* amino acid properties is defined as follows:

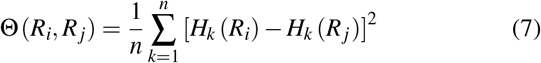

where *H_k_*(*R_i_*) denotes the *k*-th amino acid property of *R_i_*. Fourth, a set of sequence order-correlated factors is defined as follows:

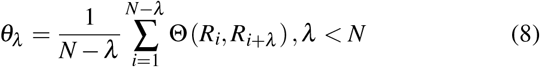

where *λ* denotes a positive integer and *N* is the integer used to define the maximum value of λ. Finally, the PAAC feature for a protein sequence is defined as follows:

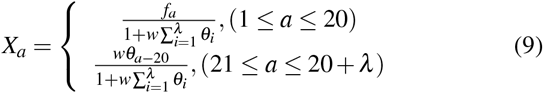

where *f_a_* denotes the normalized frequency of occurrence of amino acid *a* in the protein sequence, and w denotes the weighting factor for the sequence-order effect.

##### Quasi-Sequence Order (QSO)

The QSO feature utilizes two specific distance matrices to describe the occurrence probability of amino acids in a protein sequence, including the schneider-wrede physicochemical distance matrix [42] and the chemical distance matrix [43]. Specifically, the *d*-th rank sequence-order-coupling number is first defined as follows:

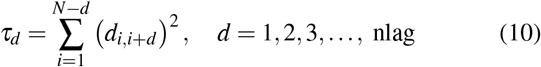

where *N* denotes the protein or peptide sequence length, *d*_*i,i*+*d*_ denotes the element in row *i*, column *i* + *d* of the distance matrix, and *nlag* denotes the maximum lag value. Then, the first 20 QSO features can be defined as follows:

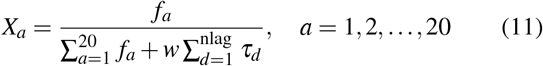

Furthermore, the other 30 QSO features are defined as:

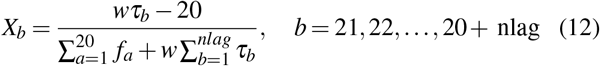

where *f_a_* denotes the normalized frequency of occurrence of amino acid *a* in the protein sequence and w denotes the weighting factor.

#### 1.2.4 Evolutionary information-based features

The Position-Specific Scoring Matrix (PSSM) consists of a set of probability scores for each amino acid (or gap) at each position in the alignment table and is used to estimate the evolutionary conservation of genes. The basic idea of PSSM is to match query sequences in a database to sequences in an alignment table, giving higher weights to conserved positions than to variable positions. In recent years, the PSSM profiles have been successfully used in various fields, including identifying functional residues, binding residues and proteins of different fold types, etc. In this work, we generated the original PSSM profiles by utilizing the PSI-BLAST program (version blast-2.12.0) [20] to iteratively (3 times) search distantly related homologous sequence of proteins against the database UniRef50 [44] with a specified e-value score (0.001).

##### PSSM-composition

The PSSM-composition feature [45] eliminates the variability introduced by protein sequence length by summing and averaging all rows of the original PSSM profile for each naturally occurring amino acid type, and is defined as follows:

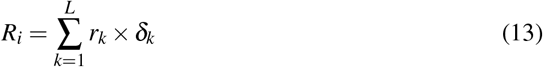

subject to

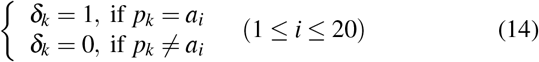

where *R_i_* denotes the *i*-th row of the PSSM-composite feature matrix, *r_k_* denotes the *k*-th row of the normalized PSSM profile, *p_k_* denotes the *k*-th amino acid in the protein sequence, and *a_i_* denotes the *i*-th amino acid of the 20 standard amino acids.

##### RPM-PSSM

The RPM-PSSM feature [46] transforms the original PSSM by filtering the negative values to 0 and leaving the positive values unchanged. The idea of the method is derived from the residue probing method, where each amino acid corresponding to a specific column in the PSSM is considered as a probe. Ultimately, the original PSSM is transformed into a 400-dimensional feature vector, using defined as follows:

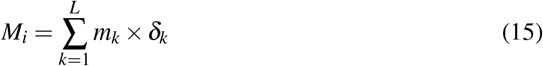

subject to

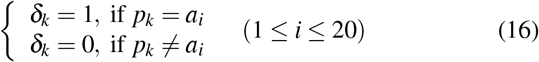

where *M_i_* denotes the *i*-th row of the RPM-PSSM feature matrix, *m_k_* denotes the *k*-th row of the original PSSM profile, *p_k_* denotes the *k*-th amino acid in the protein sequence, and *a_i_* denotes the *i*-th amino acid of the 20 standard amino acids.

##### AADP-PSSM

The AADP-PSSM [47] feature extends the traditional AAC and DPC concepts to PSSM. Firstly, the AAC-PSSM is transformed into a fixed-length 20-dimensional feature vector by averaging the columns of the original PSSM profile, defined as follows:

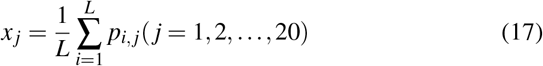

where *x_j_* denotes the *j*-th row of the AAC-PSSM feature matrix, representing the average proportion of amino acid mutations in the evolutionary process, *p_i,j_* denotes the entity in row *i* and column *j* of the original PSSM profile. Secondly, the DPC-PSSM is transformed into a fixed-length 400-dimensional feature vector to avoid information loss due to X in the protein, defined as follows:

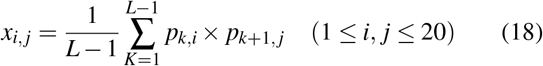

Finally, the AADP-PSSM was transformed into a fixed-length 20+400=420-dimensional feature vector by combining these two components.

#### 1.2.5 One-hot encoding feature

Unlike classical machine learning methods that require large amounts of expert knowledge to characterise genetic features prior to training, deep learning is an end-to-end model that automatically learns genetic features from large amounts of sequence data. The one-hot encoding [48] feature conveniently converts protein sequences into numerical vectors as inputs for deep learning, where each of the 20 natural amino acids is converted into a 20-dimensional feature vector, with the amino acid locations set to 1 and the remaining locations set to 0 in alphabetical order. In addition, the gene sequence lengths were intercepted uniformly at 2,000, as the distribution of gene sequence lengths in our dataset mostly falls within this range. Finally, the one-hot encoding feature is transformed into a 20×2000=40,000-dimensional feature vector.

### 1.3 Hybrid Model Training and Stacking

To obtain superior predictive performance for ARGs and VFs, HyperVR ensembles the power of classical machine learning methods and deep learning in a stacking strategy (see Figure 1.C). Stacking is an ensemble learning technique that has been proven to have better prediction results in several fields and is first recommended in many high-level competitions [49, 50]. It integrates multiple base-level classification or regression models through a single meta-classifier or meta-regressor. The base-level model does the training using the entire training dataset, and the meta-model uses the output of the base-level models as features for training. To address the over-fitting phenomenon in the final prediction, we further utilized the 5-fold cross-validation method to respectively train the base-level models. The detailed training process is shown in Supplementary File Figure S1.

In particular, classical machine learning methods in HyperVR are mainly chosen to produce better predictive performance with ensemble learning methods, including Random Forest [51], Extra Trees classifier [52], Xgboost [53], GradientBoosting [54] and Adaboost [55]. The stacking algorithm in HyperVR is represented by the pseudocode shown in Algorithm 1.

#### Algorithm 1 Stacking in HyperVR

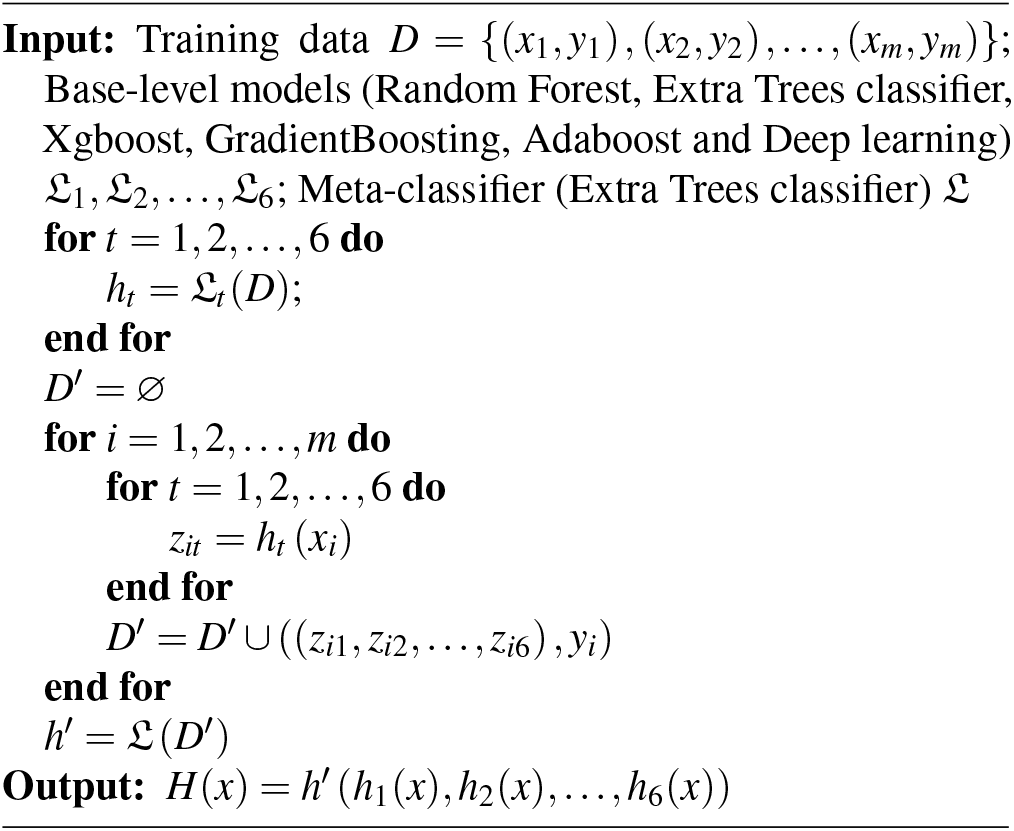

In detail, for random forest and extra trees classifier, the sklearn package in python is adopted, the number of trees in the forest is set to 500, the information gain method is selected to Gini impurity, and the maximum depth of the trees is set to none. The standalone xgboost package in python is adopted, and the maximum number of estimators to terminate the boost is set to 500, the booster is selected to GBTree, and the other parameters are set to their default values. For adaboost and gradient boosting, the sklearn package in python is also adopted and the maximum number of estimators to terminate the boost is set to 500. In addition, the learning rate and booster of adaboost is respectively set to 1.0 and SAMME.R. The loss function to be optimized and the function to measure the quality of a split in gradient boosting is respectively set to Deviance and Friedman_mse.

In this work, deep learning is used to target automatically training two classes of the bit score-based similarity and one-hot encoding features that do not require prior knowledge. Specifically, we construct the deep learning framework containing six hidden layers, where the number of neurons is respectively (2^12^, 2^10^, 2^8^, 2^6^, 2^4^, 2^2^). Furthermore, a dropout layer follows each hidden layer to avoid over-fitting, and the dropout rate is set to 0.05. The model ends with the sigmoid layer, which is used to output the final predicted scores. In addition, the framework is written by the TensorFlow program and compiled with the following parameters, where the optimizer is the stochastic gradient descent (SGD) method with a learning rate of 0.05, the loss function is BinaryCrossentropy, the evaluation metric is BinaryAccuracy, the training epoch is 500 and early stopping mechanism is added.

### 1.4 Evaluation Criteria

In this work, HyperVR was evaluated using the commonly multi-label standard performance metric, including precision, recall, F1-score, and their micro-averages. The specific formulas are defined as follows:

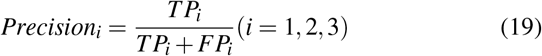

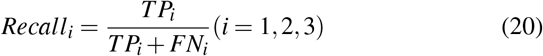

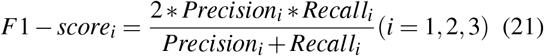

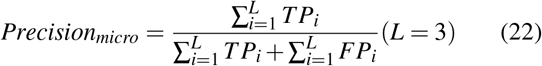

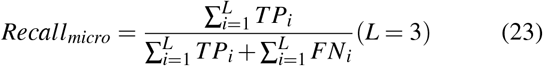

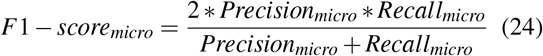

where TP (true positive), FP (false positive), TN (true negative), and FN (false negative) respectively denote the number of positive samples correctly labeled, negative samples incorrectly labeled, negative samples correctly labeled, and positive samples incorrectly labeled. The *i* = 1,2,3 respectively represents the ARG, VF, and NS category, and *L* = 3 is the total number of categories.

## 2. Results and Discussion

### 2.1 Implementation Details

The classic machine learning model and deep learning network of HyperVR were respectively implemented on Python 3.8 with Scikit-learn 1.0.1 and Keras 2.8.0 with Tensorflow-gpu 2.8.0 backend. In addition, we utilized Keras multi-GPU processing to increase the training speed significantly. The experiments were performed on a Linux system server with 16x Intel(R) Xeon(R) Bronze 3106 CPUs (1.70GHz) featuring 128 CPU cores in total, 4x NVIDIA V100 GPUs, and 250GB of RAM.

### 2.2 HyperVR can accurately predict ARGs, VFs, and NSs simultaneously

To accurately evaluate the performance of HyperVR in predicting ARGs, VFs, and NSs simultaneously, the 5-fold crossvalidation method was employed in this section, resulting in an average accuracy of 91.94%. We implemented the rigorous procedure that cross-validation step to enable an unbiased evaluation of the effectiveness of HyperVR. After training HyperVR with 80% of the selected raw data from HyperVR-DB, the remaining 20% held-out data was utilized to evaluate its generalization capabilities, and so on for five trials repeated (Figure 2.A).

**Figure 2.**
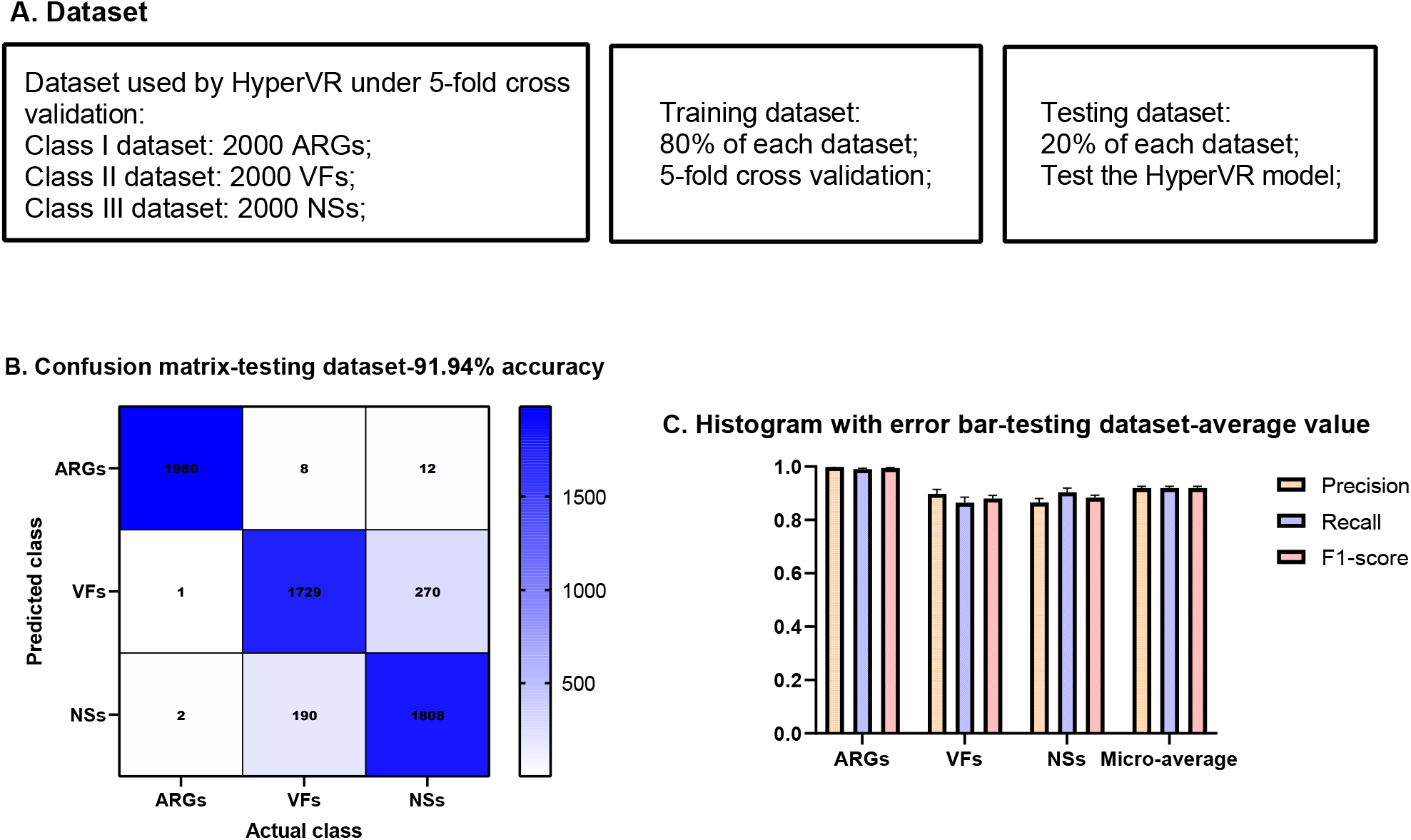
The prediction results of HyperVR to simultaneously predict ARGs, VFs, and NSs under 5-fold cross-validation. **A**. The dataset used by HyperVR under 5-fold cross validation. **B**. The confusion matrix generated by HyperVR under 5-fold cross validation. **C**. The histogram with error bar of evaluation metrics under 5-fold cross validation.

We recorded detailed classification reports, including accuracy, precision, recall, F1-score, micro-average metrics (Supplementary File Table S2), and plotted confusion matrices (Supplementary File Figure S2) for each fold. The mean and standard deviation of all metrics derived from all 5 cross-validation experiments were calculated and reported in Table 1. Figures 2.B and 2.C respectively visualize the final confusion matrix generated in the experiment and the error bar histograms of three evaluation indicators. All the results reported above demonstrate that HyperVR can accurately classify ARGs, VFs, and NSs at the same time, especially with the excellent performance of 99.85% precision and 99% recall for ARGs, and 89.76% accuracy and 86.45% recall for VFs. Furthermore, the standard deviation of 0.65% for the micro-average results under five independent experiments further demonstrates the stability of HyperVR.

**Table 1.**
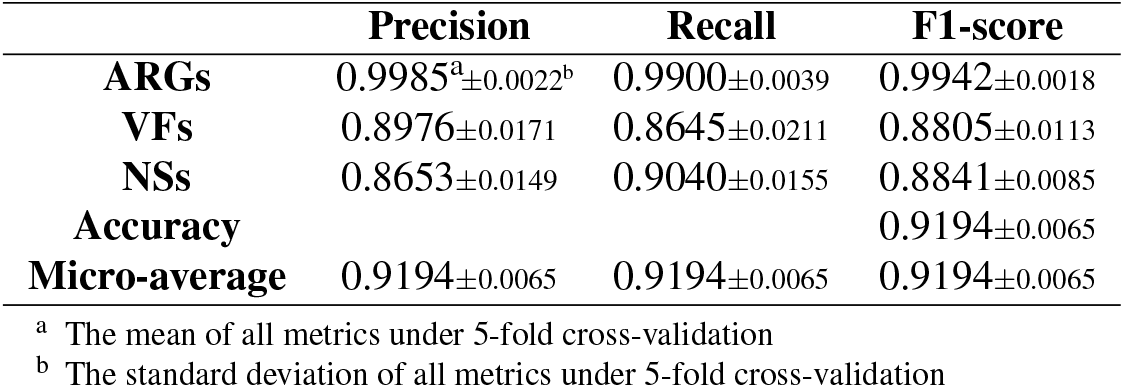
The detailed results of HyperVR to simultaneously predict ARGs, VFs, and NSs under 5-fold cross-validation

|  | Precision | Recall | F1-score |
| --- | --- | --- | --- |
| **ARGs** | 0.9985<sup>a</sup>±0.0022<sup>b</sup> | 0.9900±0.0039 | 0.9942±0.0018 |
| **VFs** | 0.8976±0.0171 | 0.8645±0.0211 | 0.8805±0.0113 |
| **NSs** | 0.8653±0.0149 | 0.9040±0.0155 | 0.8841±0.0085 |
| **Accuracy** |  |  | 0.9194±0.0065 |
| **Micro-average** | 0.9194±0.0065 | 0.9194±0.0065 | 0.9194±0.0065 |

<sup>a</sup> The mean of all metrics under 5-fold cross-validation<sup>b</sup> The standard deviation of all metrics under 5-fold cross-validation

### 2.3 HyperVR-ARGs can flexibly and accurately predict ARGs individually

To further evaluate the performance of HyperVR in predicting ARGs individually (shorthand HyperVR-ARGs), the same 5-fold cross-validation method was employed in this section, resulting in an excellent average cross-validation accuracy of 99.85%. The detailed cross-validatione results for each part in HyperVR-ARGs, including accuracy (Acc.), sensitivity (Sen.), specificity (Spec.), precision (Prec.), matthews correlation coefficient (MCC), and the areas under the ROC curve (AUC), were recorded in Supplementary File Table S3. The final results of HyperVR-ARGs under 5-fold cross-validation were shown in Table 2. On the other hand, we plotted the receiver operating characteristic (ROC) curve (Figure 3.A) and the precision-recall (PR) curve (Figure 3.B) to visualize the performance of HyperVR-ARGs. Figure 3.C visualized the training process of the DNN for the Bit score-based similarity feature. Figure 3.D further visualized the prediction results of each part in HyperVR-ARGs under 5-fold cross-validation through the error bar histogram.

**Table 2.** The 5-fold cross-validation results of HyperVR-ARGs to predict ARGs individually

| Fold | Acc. | Sen. | Spec. | Prec. | MCC | AUC |
| --- | --- | --- | --- | --- | --- | --- |
| 1st | 0.9967 | 0.9950 | 0.9975 | 0.9950 | 0.9925 | 0.9985 |
| 2nd | 0.9958 | 0.9925 | 0.9975 | 0.9950 | 0.9906 | 0.9984 |
| 3rd | 0.9958 | 0.9875 | 1.0000 | 1.0000 | 0.9906 | 0.9984 |
| 4th | 0.9967 | 0.9950 | 0.9975 | 0.9950 | 0.9925 | 0.9985 |
| 5th | 0.9942 | 0.9825 | 1.0000 | 1.0000 | 0.9869 | 0.9984 |
| Average±s.t.d | 0.9958±0.0010 | 0.9905±0.0054 | 0.9985±0.0014 | 0.9970±0.0027 | 0.9906±0.0023 | 0.9984±0.0001 |

**Figure 3.**
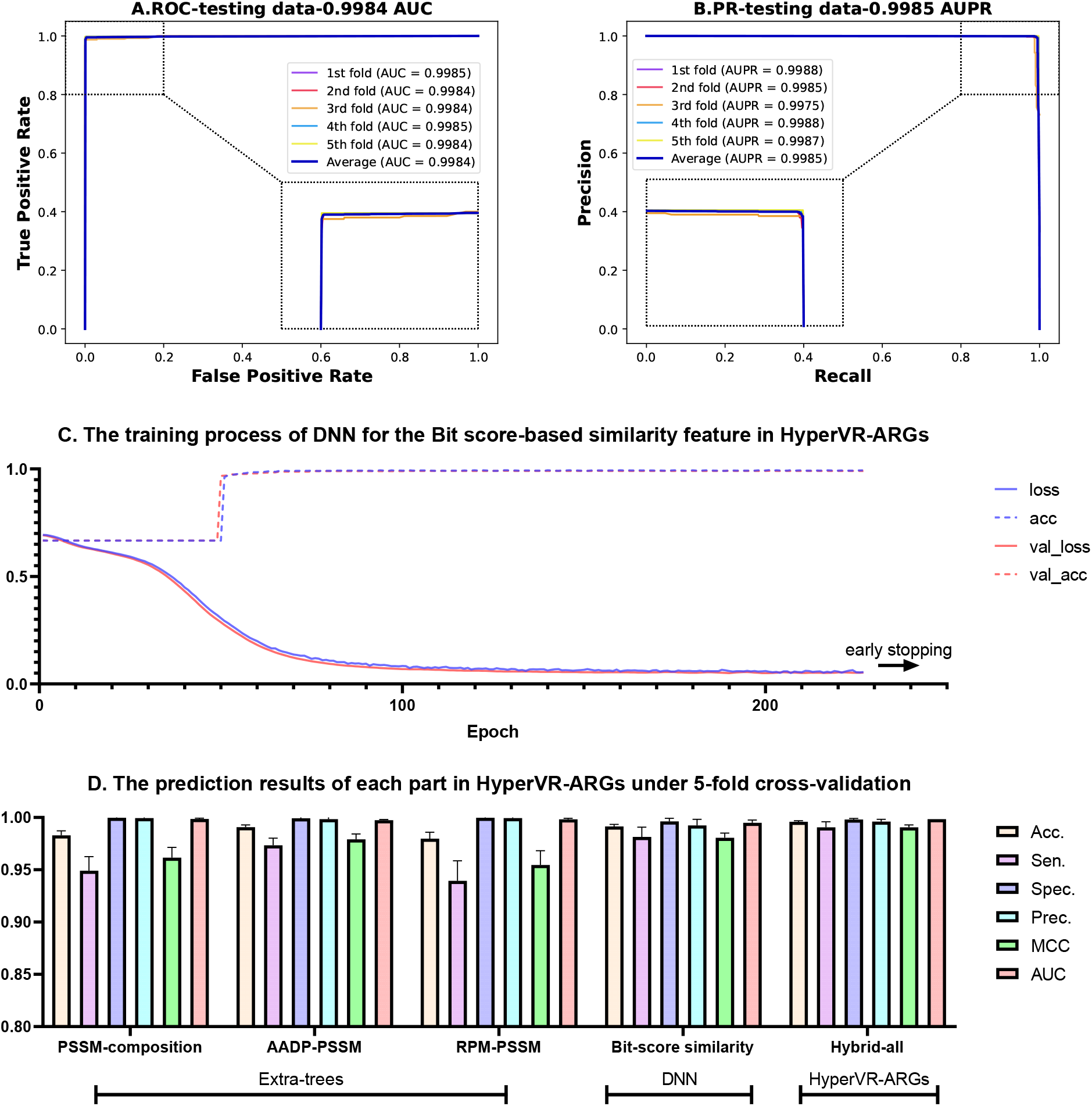
The 5-fold cross-validation performance for HyperVR-ARGs to predict ARGs individually **A**. The ROC curves and interpolated area under ROC curve (AUC) of HyperVR-ARGs under 5-fold cross validation. **B**. The PR curves and interpolated area under PR curve (AUPR) of HyperVR-ARGs under 5-fold cross validation. **C**. The training process of DNN for the Bit score-based similarity feature in HyperVR-ARGs. **D**. The prediction results of each part in HyperVR-ARGs under 5-fold cross validation.

The predictive power of HyperVR-ARGs comes from two contributions, including traditional machine learning methods (Extra-trees classifier) utilizing gene evolutionary information in the form of PSSM and DNN utilizing Bitscore-based similarity features of full gene sequences. From the results in this section, we can first see that the DNN can quickly reach high accuracy (about 50 epochs) and converge quickly with the early stopping mechanism, which tells us that using the combination of DNN and Bit-score based similarity feature is fast and effective for identifying ARGs. Secondly, the comparison results in the Figure 3.D show that excellent performance in identifying ARGs can also be achieved using traditional machine learning methods and the corresponding PSSM features, especially the combination of Extra-trees and RPM-PSSM. Finally, HyperVR-ARGs can achieve better predictive performance by combining the capabilities of different combinations. In summary, HyperVR-ARGs can flexibly and accurately predict ARGs individually.

### 2.4 HyperVR-VFs can flexibly and accurately predict VFs individually

To further estimate the predictive ability of HyperVR in predicting VFs individually (shorthand HyperVR-VFs), the 5-fold cross-validation in this section was performed again, resulting in a good average cross-validation accuracy of 91.83%. The same evaluation strategy as in Section 2.3 was adopted. The detailed cross-validation results for each part in HyperVR-VFs were shown in Supplementary File Table S4. The final stacking results of HyperVR-VFs under 5-fold crossvalidation were shown in Table 3. We also calculated the AUC of the ROC curves and PR curves of HyperVR-VFs under 5-fold cross-validation, as shown in Figures 4.A and B. Figure 4.C illustrated the training process of the DNN for the one-hot encoding feature. Figure 4.D visualized the prediction AUC results of each part in HyperVR-VFs under 5-fold cross-validation through the error bar histogram.

**Table 3.**
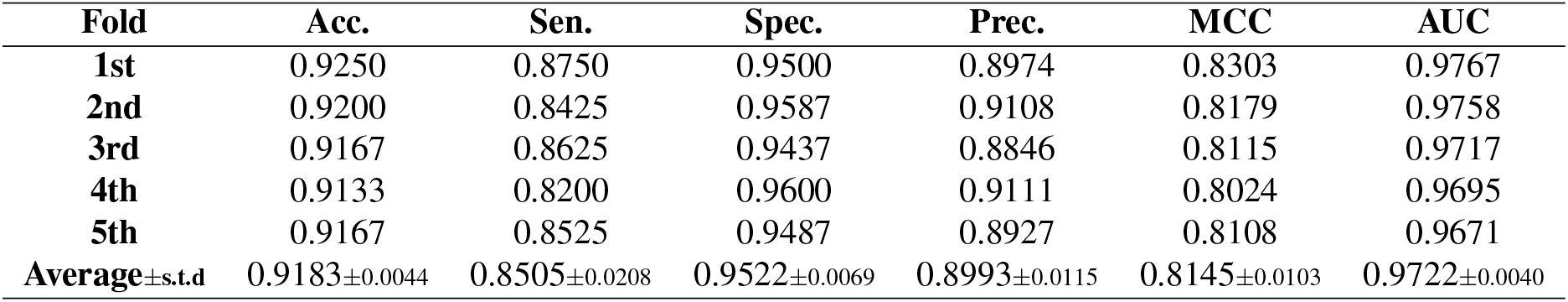
The 5-fold cross-validation results of HyperVR-VFs to predict VFs individually

**Figure 4.**
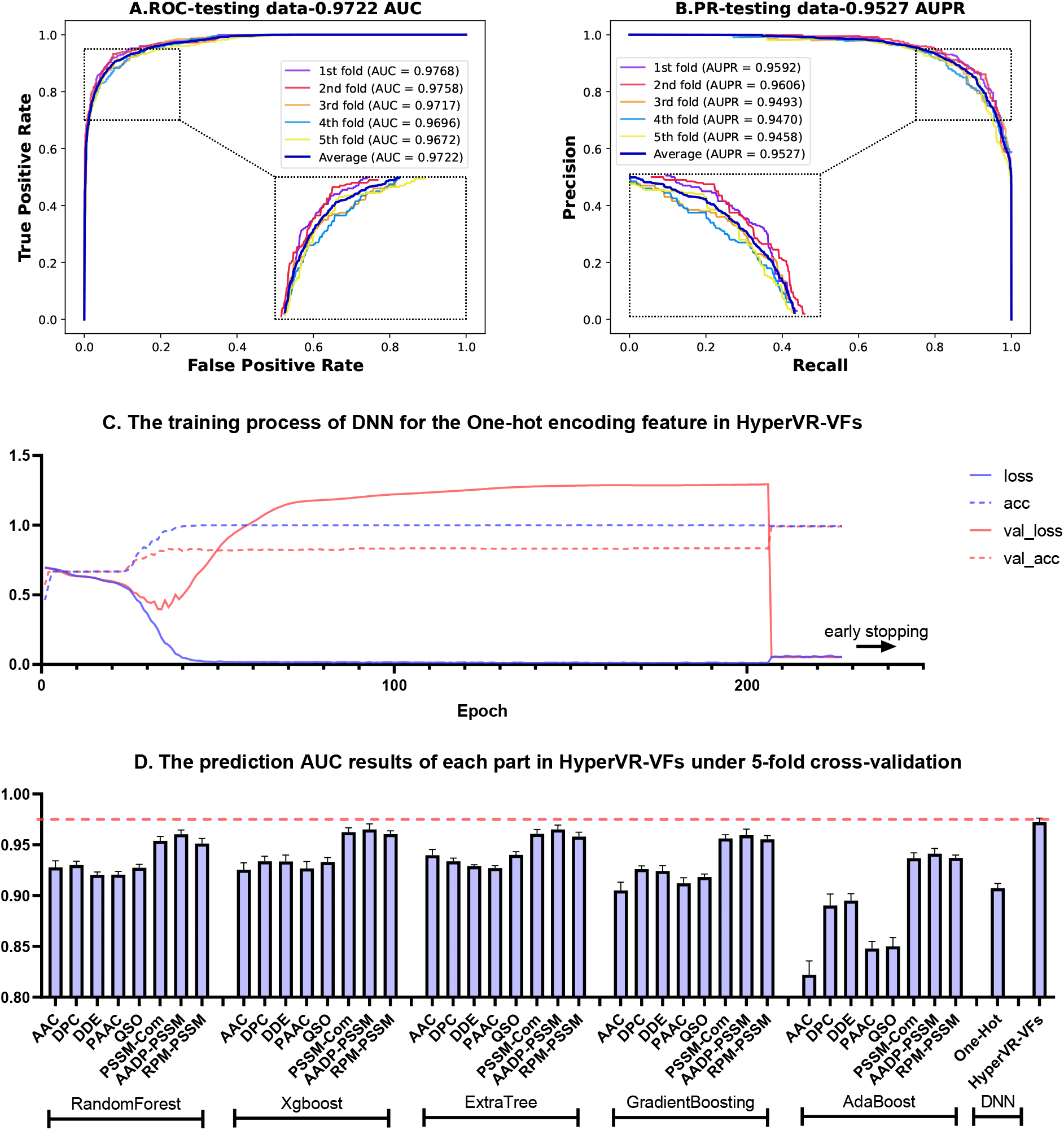
The 5-fold cross-validation performance for HyperVR-VFs to predict VFs individually **A**. The ROC curves and interpolated area under ROC curve (AUC) of HyperVR-VFs under 5-fold cross validation. **B**. The PR curves and interpolated area under PR curve (AUPR) of HyperVR-VFs under 5-fold cross validation. **C**. The training process of DNN for the One-hot encoding feature in HyperVR-VFs. **D**. The prediction AUC results of each part in HyperVR-VFs under 5-fold cross validation.

The predictive power of HyperVR-VFs comes from more contributions, including traditional ensemble learning methods utilizing sequence-based features, physicochemical property-based features, and gene evolutionary information in the form of the PSSM and DNN utilizing One-hot encode features. From the results in this section, we can see that the DNN reached best performance and converge on the validation dataset after about 210 epochs with the early stopping mechanism. The evolutionary information of genes contribute more than sequence-based features, physicochemical property-based features and One-hot encode features for the prediction of VFs. Last but not least, the stacking of these multiple combinations can yield better prediction performance than any one of them. It can be concluded that HyperVR-VFs have a strong and stable ability to predict VFs individually.

### 2.5 Validation of HyperVR through novel ARGs, VFs and NSs

To test the predictive ability of HyperVR on novel ARGs, VFs and NSs, an independent dataset in HyperVR-DB was obtained consisting of 209 ARGs, 209 VFs and 209 NSs. Notably, the novel genes were completely independent from the genes in the training dataset, removing all identical or duplicate sequences by setting the identity threshold to 100% of CD-HIT. Moreover, we also introduced the the currently available VRprofile [25] model and the traditional best hit approach as baseline comparison methods. VRprofile is used to find virulence and/or drug resistance genes and their transfer-associated gene clusters by quickly performing homology searches from user-supplied pathogenic bacterial genome sequences. The best hit approach is presently the most commonly used methods for identifying gene classes based on the computational principle of comparing the gene sequences against available online databases. Such comparison aligned raw reads or predicted open reading frames (full gene length sequences) from assembled contigs to the database of choice by using programs such as BLAST [56], Bowtie [22], or DIAMOND [21], and then predicting or assigning the categories of genes to present using a sequence similarity cutoff or sometimes an alignment length requirement. Since both methods require a choice of thresholds, we choose three different thresholds for the experiments respectively, and compare their experimental results with those of HyperVR. Specifically, we chose Ha-value parameters of 0.21, 0.64, and 0.81 for VRprofile, according to their integrated Web interface (https://bioinfo-mml.sjtu.edu.cn/VRprofile/). For the best hit approach, we utilized the DIAMOND program and chose the identity parameters of 0.21 (Diamond-21%), 0.64 (Diamond-64%) and 0.81 (Diamond-81%) for consistency.

Table 4 shows the detailed prediction results of HyperVR and two different baseline methods under different parameters for these novel ARGs, VFs, and NSs. It should be noted that we find that VRprofile obtains the same prediction results for three different Ha-value parameters, so we only list one of the results. Figure 5 visualized the confusion matrices and histograms of results for different baseline methods. Thus, we can draw the following tentative conclusions from the foregoing results. VRprofile has good precision and recall for identifying novel ARGs, but very poor identification of VFs. The best hit approach using the DIAMOND program is sensitive to the identity cutoffs. When the identity cutoff is relatively small, the best hit approach is relatively good at identifying ARGs but still poor at identifying VFs. By increasing the identity cutoff, the identification precision of ARGs and VFs will be improved, but at the same time the recall will be greatly reduced. Overall, the best hit approach does not yield an overall superior prediction for the novel ARGs and VFs. In contrast, HyperVR achieved excellent overall performance in predicting both novel ARGs and VFs simultaneously without parameter selection. This again proves that HyperVR can be used as an effective and simple tool for the identification of ARGs and VFs.

**Table 4.** The prediction results of HyperVR and baseline methods for novel ARGs, VFs, and NSs

|  |  | Precision | Recall | F1-score |  |  | Precision | Recall | F1-score |
| --- | --- | --- | --- | --- | --- | --- | --- | --- | --- |
| VRprofile | ARGs | 0.9801 | 0.9426 | 0.9610 | Diamond-21% | ARGs | 0.9533 | 0.9761 | 0.9645 |
|  | VFs | 0.5490 | 0.1340 | 0.2154 |  | VFs | 0.5189 | 0.4593 | 0.4873 |
|  | NSs | 0.5227 | 0.9378 | 0.6712 |  | NSs | 0.5175 | 0.5646 | 0.5400 |
|  | Accuracy |  |  | 0.6715 |  | Accuracy |  |  | 0.6667 |
|  | Mico-average | 0.6715 | 0.6715 | 0.6715 |  | Mico-average | 0.6667 | 0.6667 | 0.6667 |
|  |  | Precision | Recall | F1-score |  |  | Precision | Recall | F1-score |
| Diamond-64% | ARGs | 1.0000 | 0.7129 | 0.8324 | Diamond-81% | ARGs | 1.0000 | 0.4976 | 0.6645 |
|  | VFs | 0.7805 | 0.1531 | 0.2560 |  | VFs | 0.9375 | 0.1435 | 0.2490 |
|  | NSs | 0.4577 | 0.9569 | 0.6192 |  | NSs | 0.4216 | 0.9904 | 0.5914 |
|  | Accuracy |  |  | 0.6077 |  | Accuracy |  |  | 0.5439 |
|  | Mico-average | 0.6077 | 0.6077 | 0.6077 |  | Mico-average | 0.5439 | 0.5439 | 0.5439 |
|  |  | Precision | Recall | F1-score |  |  | Precision | Recall | F1-score |
| HyperVR | ARGs | 0.9952 | 1.0000 | 0.9976 |  |  |  |  |  |
|  | VFs | 0.9351 | 0.8278 | 0.8782 |  |  |  |  |  |
|  | NSs | 0.8448 | 0.9378 | 0.8889 |  |  |  |  |  |
|  | Accuracy |  |  | 0.9219 |  |  |  |  |  |
|  | Mico-average | 0.9219 | 0.9219 | 0.9219 |  |  |  |  |  |

**Figure 5.**
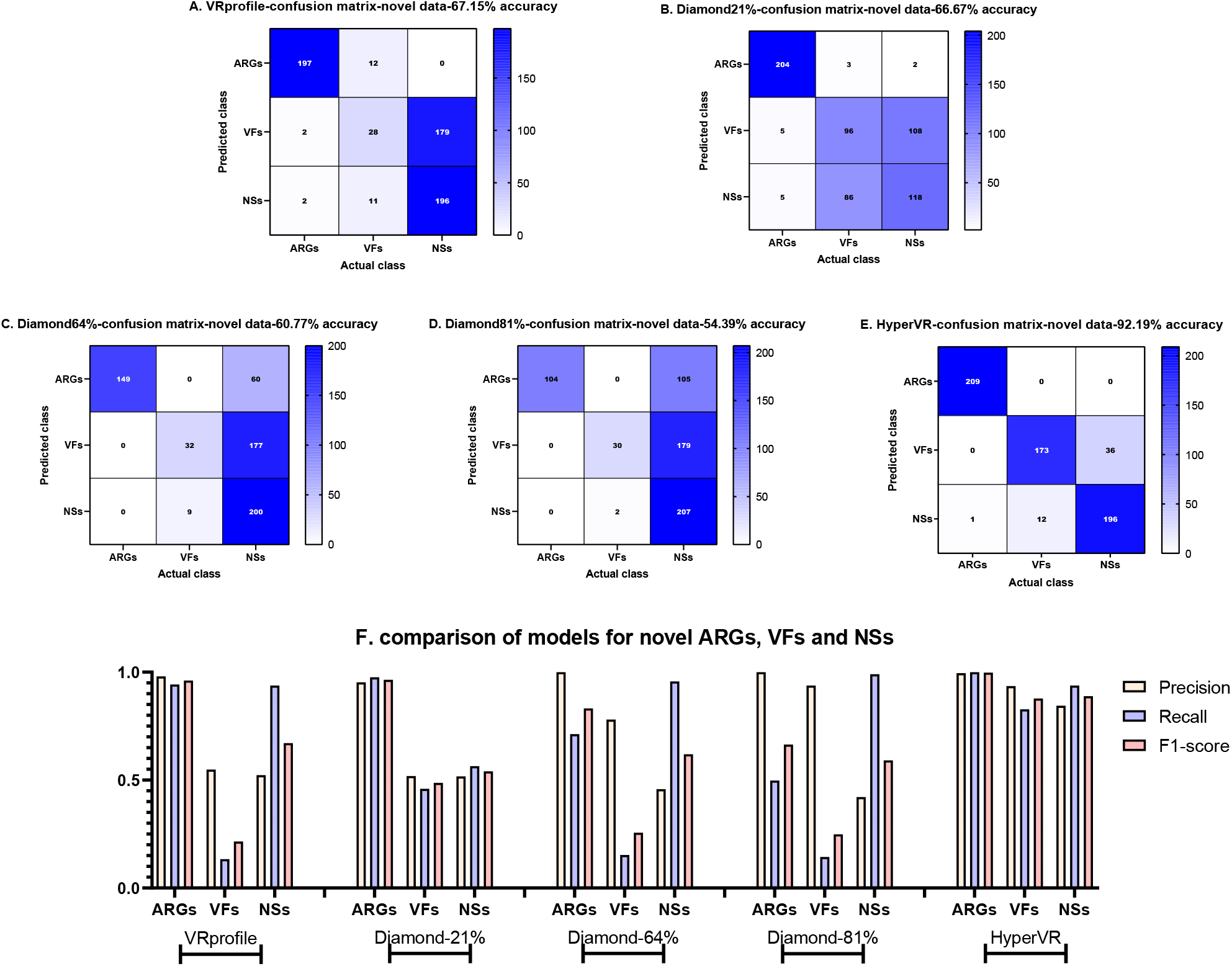
The confusion matrices and histograms representing results for HyperVR and baseline methods to simultaneously predict novel ARGs, VFs, and NSs. **A**. The confusion matrix of VRprofile for simultaneously predicting novel ARGs, VFs, and NSs. **B**. The confusion matrix of best hit approach by using DIAMOND program with 21% identity for simultaneously predicting novel ARGs, VFs, and NSs. **C**. The confusion matrix of best hit approach by using DIAMOND program with 64% identity for simultaneously predicting novel ARGs, VFs, and NSs. **D**. The confusion matrix of best hit approach by using DIAMOND program with 81% identity for simultaneously predicting novel ARGs, VFs, and NSs. **E**. The confusion matrix of HyperVR for simultaneously predicting novel ARGs, VFs, and NSs. **F**. The histograms representing results for HyperVR and baseline methods to simultaneously predict novel ARGs, VFs, and NSs.

### 2.6 Validation of HyperVR through an in Silico spikein experiment

For metagenomic datasets from real-world samples, ARGs and VFs may represent only a small fraction of the total number of genes. It is essential to evaluate the performance of HyperVR in cases where non-target genes dominate. To assess the ability of HyperVR for predicting a small number of ARGs and VFs among the majority of NSs, we constructed a negative metagenomic dataset mimicking a spike-in metagenomic experiment. Specifically, we first randomly selected 10 ARGs and 10 VFs from the independent dataset. Next, we reconstructed 10,000 negative samples using the negative sample construction method (refer to Section 1.1.3). The final spike-in metagenomics dataset containing 10020 genes were ensured to have no overlap with the training dataset, and the percentage of positive samples was only (20/10020)% ≈ 0.19%. In this section, we have chosen the same baseline methods as Section 2.5 for comparison. Table 5 shows the prediction results of HyperVR and baseline methods for positive samples in the spike-in metagenomics dataset. From the table, we see that VRprofile has a good identification effect for a small number of ARGs among the majority of NSs, but the identification effect for VFs is extremely poor. The best hit approach using the DIAMOND program decreases in identification accuracy as the identity cutoff increases, and the best result is achieved when the identity is 21%, which is better than the VRprofile method. HyperVR obtained the best prediction results among the three methods, with all 10 ARGs predicted correctly and only 1 of 10 VFs predicted incorrectly to negative sample. This section demonstrates that HyperVR can well predict simultaneously small amounts of ARGs and VFs that exist in a large number of negative samples, which is applicable to the fact that ARGs and VFs may represent only a small fraction of the total number of genes in the real world.

**Table 5.**
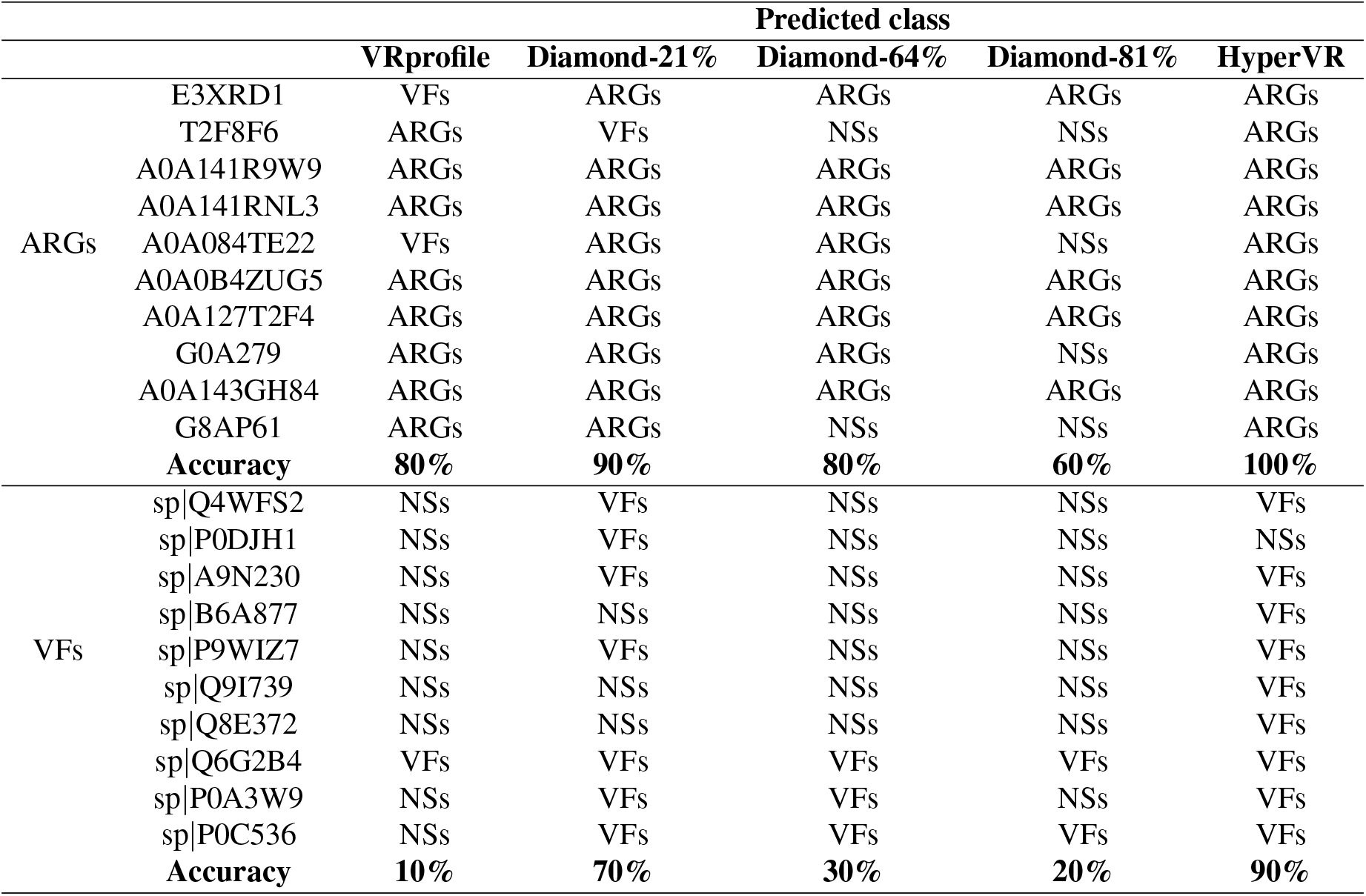
The prediction results of HyperVR and baseline methods through an in Silico spike-in experiment

### 2.7 Validation of HyperVR through Pseudo ARGs and VFs

To verify the predictive power of HyperVR for pseudo ARGs and VFs, which contain segments of ARGs or VFs but are not true ARGs or VFs, a false positive dataset containing 300 pseudo-ARGs and 300 pseudo-VFs was constructed. To construct one of these genes, we first randomly picked the k-mer of 50 amino acids long of ARGs or VFs from our independent dataset, respectively. Next, five k-mers of 50 amino acids long were randomly picked from negative samples in our independent dataset and randomly spliced with the ARGs and VFs segments generated in the first step. For comparison, the baseline methods in Sections 2.5 and 2.6 are utilized again. In the results, VRprofile misidentified 3 genes as ARGs, with a false positive rate of (3/600=0.50%). In addition, Diamond-21% misidentified 435 genes including 298 ARGs and 137 VFs, with a false positive rate of (435/600=72.50%), Diamond-64% misidentified 320 genes including 244 ARGs and 76 VFs, with a false positive rate of (320/600≈53.33%), Diamond-81% misidentified 219 genes including 176 ARGs and 43 VFs, with a false positive rate of (219/600=36.50%), and HyperVR misidentified 66 genes including 31 ARGs and 35 VFs, with a false positive rate of (66/600=11%). Although HyperVR has a higher false positive rate than VRprofile, considering their performance in Sections 2.5 and 2.6, the overall predictive power of HyperVR is still better, and certainly better than the best hit approach. This section further demonstrates that the ability of HyperVR to simultaneously predict ARGs and VFs does not just memorize the gene sequences of the training samples, but actually learns the uniqueness of ARGs and VFs by considering multiple features and classifiers together.

### 2.8 Validation of HyperVR through pathogens cases

To test the ability of HyperVR to simultaneously identify ARGs and VFs in real pathogenic bacteria, three specific pathogen strain datasets were chosen, including Mycobacterium tuberculosis (strain ATCC 25618 / H37Rv, 3993 protein genes) [57], Bacillus anthracis (strain Ames Ancestor, 5493 protein genes) [58] and Staphylococcus aureus (strain NCTC 8325 / PS 47, 2889 protein genes) [59]. All three of these pathogens are focus of current research. Specifically, tuberculosis is, to this day, according to the WHO, the leading killer of adults, with approximately 2 million deaths annually worldwide. An additional problem for tuberculosis is the emergence of drug resistant strains, mainly because people do not complete their treatment plans or have been incorrectly treated and so have remained infectious [57]. Bacillus anthracis is an endospore-forming bacterium that causes inhalational anthrax [58]. Staphylococcus aureus is one of the major causes of community-acquired and hospital-acquired infections [59].

Each of the three strain datasets was respectively input to HyperVR as a test dataset for simultaneous identification of ARGs and VFs. We listed the genes that HyperVR considered most likely to be ARGs or VFs (prediction scores >95%, increase the threshold if a high-quality set of ARGs and VFs prediction is desired). Subsequently, we verified each of these genes in the following two ways. Firstly, the genes were directly queried in existing databases and determined by database annotation. Secondly, blast was used to find similar genes and the type of genes was determined by database annotation of similar genes. The genes predicted by HyperVR in Mycobacterium tuberculosis, Bacillus anthracis and Staphylococcus aureus are respectively shown in Supplementary File Table S5, S6 and S7. It is evident from the results that HyperVR can effectively identify ARGs and VFs in real pathogenic strains, which narrows down the scope for researchers to conduct in vitro experiments and greatly saves them time and effort. Furthermore, it should be noted that, as with all in silico predictions, HyperVR is used to obtain an overview or inference of ARGs and VFs in a pathogenic strain. Some genes are not explicitly annotated as antibiotic resistant or virluence by the database, but it may still be the corresponding gene, e.g. genes annotated as penicillin binding, toxin etc. Strictly speaking, downstream experiments are required to determined which category the gene truly belongs to.

## 3. Conclusions

Here, a novel hybrid prediction framework called HyperVR for simultaneously predicting virulence factors and antibiotic resistance genes was proposed. To the best of our knowledge, this is the first computational tool that can simultaneously predict ARGs and VFs. HyperVR works by integrating multiple key genetic features, including bit score-based similarity feature, physicochemical property-based features, evolutionary information-based features and one-hot encoding feature, and stacking the power of classical ensemble learning methods and deep learning. It can accurately predict VFs, ARGs and negative samples at the same time, and can be used flexibly and accurately to predict VFs or ARGs individually. In addition, HyperVR addresses the drawbacks of traditional best-hit methods with high false negatives and the need for strict cut-off thresholds, while outperforming previous tools in terms of precision and recall on novel VFs and ARGs, silico spike-in VFs and ARGs, and pseudo VFs and ARGs. Overall, HyperVR is an effective, simple prediction tool and requires only gene sequence information without additional expert knowledge input.

However, HyperVR also has some limitations that we need to further optimize in the future. For example, HyperVR uses multiple features and models for training, resulting in a longer training time for the model, but the good thing is that the training process only needs to be performed once. Second, HyperVR is not as accurate as ARGs for predicting VFs, mainly because the bit score-based similarity feature does not distinguish well between VFs and NSs, which is consistent with the poor identification of VFs by traditional best-hit methods and inspires us to focus on improving the prediction accuracy of VFs. Furthermore, HyperVR currently only predicts whether genes are VFs or ARGs, or neither, and cannot accurately predict the specific class of genes. This will also be the next step we need to address.

## Acknowledgments

This work was supported by National Key R&D Program of China 2017YFB0202602, 2018YFC0910405, 2017YFC1311003, 2016YFC1302500, 2016YFB0200400, 2017YFB0202104; NSFC Grants U19A2067, 61772543, U1435222, 61625202, 61272056, 62102427, 61762031; Science Foundation for Distinguished Young Scholars of Hunan Province (2020JJ2009); Science Foundation of Changsha kq2004010; JZ20195242029, JH20199142034, Z202069420652; The Funds of Peng Cheng Lab, State Key Laboratory of Chemo/Biosensing and Chemometrics; the Fundamental Research Funds for the Central Universities, and Guangdong Provincial Department of Science and Technology under grant No.2016B090918122.

