## Supplemental for "HyperVR–A hybrid prediction framework for virulence factors and antibiotic resistance genes in microbial data"

Infectious diseases, particularly bacterial infections, are emerging at an unprecedented rate, posing a serious challenge to public health and the global economy. Different virulence factors (VFs) work in concert to enable pathogenic bacteria to successfully adhere, reproduce and cause damage to host cells, and antibiotic resistance genes (ARGs) allow pathogens to evade otherwise curable treatments. To understand the causal relationship between microbiome composition, function and disease, both VFs and ARGs in microbial data must be identified. Most existing computational models cannot simultaneously identify VFs or ARGs, hindering the related research. The best hit approaches are currently the main tools to identify VFs and ARGs concurrently; yet they usually have high false-negative rates and are very sensitive to the cut-off thresholds. In this work, we proposed a hybrid computational framework called HyperVR to predict VFs and ARGs at the same time. Specifically, HyperVR integrates key genetic features and then stacks classical ensemble learning methods and deep learning for training and prediction. HyperVR accurately predicts VFs, ARGs and negative genes (neither VFs nor ARGs) simultaneously, with both high precision (>0.91) and recall (>0.91) rates. Also, HyperVR keeps the flexibility to predict VFs or ARGs individually. Regarding novel VFs and ARGs, the VFs and ARGs in metagenomic data, and pseudo VFs and ARGs (gene fragments), HyperVR has shown good prediction, outperforming the current state-of-the-art predition tools and best hit approaches in terms of precision and recall. HyperVR is a powerful tool for predicting VFs and ARGs simultaneously by using only gene sequences and without strict cut-off thresholds, hence making prediction straightforward and accurate.

### SUPPLEMENTARY MATERIAL

### TABLE OF CONTENTS

#### A. Supplementary Tables

Table S1. The removed genes containing the virulence or antibiotic keywords in the UNIPROT Swiss-Prot database.
Table S2. The detailed classification report of HyperVR under 5-fold cross-validation.
Table S3. The detailed cross-validation results for each part in HyperVR-ARGs under 5-fold cross validation.
Table S4. The detailed cross-validation results for each part in HyperVR-VFs under 5-fold cross validation.
Table S5. The genes predicted by HyperVR with >95% score in Mycobacterium tuberculosis strains.
Table **??**. The genes predicted by HyperVR with >95% score in Bacillus anthracis strains.
Table **??**. The genes predicted by HyperVR with >95% score in Staphylococcus aureus strains.

#### B. Supplementary Figures

Figure S1. The detailed process of HyperVR using a stacking strategy.
Figure S2. The detailed confusion matrices of HyperVR under 5-fold cross validation.
Figure S3. The detailed confusion matrices of HyperVR-ARGs under 5-fold cross validation.
Figure S4. The detailed confusion matrices of HyperVR-VFs under 5-fold cross validation.

### SUPPLEMENTARY TABLES

**Table S1.** The removed genes containing the virulence or antibiotic keywords in the UNIPROT Swiss-Prot database

| | KeyWords | Name | Number |
|---|---|---|---|
| | KW-0045 | Antibiotic biosynthesis | 612 |
| **ARGs** | KW-0046 | Antibiotic resistance | 2280 |
| | KW-0051 | Antiviral defense | 911 |
| | KW-0843 | Virulence | 4085 |
| | KW-0945 | Host-virus interaction | 6069 |
| | KW-1160 | Virus entry into host cell | 2605 |
| | KW-1180 | Syncytium formation induced by viral infection | 21 |
| **VFs** | KW-1188 | Viral release from host cell | 931 |
| | KW-1194 | Viral DNA replication | 216 |
| | KW-1195 | Viral transcription | 236 |
| | KW-1250 | Viral genome excision | 19 |
| | KW-1251 | Viral latency | 47 |
| | KW-1272 | Viral reactivation from latency | 36 |
| | KW-1277 | Toxin-antitoxin system | 655 |

**Table S2.** The detailed classification report of HyperVR under 5-fold cross validation

| | | Precision | Recall | F1-score |
|---|---|---|---|---|
| **1st fold** | **ARGs** | 1.0000 | 0.9925 | 0.9962 |
| | **VFs** | 0.9070 | 0.8775 | 0.8920 |
| | **NSs** | 0.8774 | 0.9125 | 0.8946 |
| | **Accuracy** | | | 0.9275 |
| | **Micro-average** | 0.9275 | 0.9275 | 0.9275 |
| | | **Precision** | **Recall** | **F1-score** |
| **2nd fold** | **ARGs** | 0.9950 | 0.9900 | 0.9925 |
| | **VFs** | 0.9156 | 0.8675 | 0.8909 |
| | **NSs** | 0.8676 | 0.9175 | 0.8919 |
| | **Accuracy** | | | 0.9250 |
| | **Micro-average** | 0.9250 | 0.9250 | 0.9250 |
| | | **Precision** | **Recall** | **F1-score** |
| **3rd fold** | **ARGs** | 1.0000 | 0.9875 | 0.9937 |
| | **VFs** | 0.8784 | 0.8850 | 0.8817 |
| | **NSs** | 0.8781 | 0.8825 | 0.8803 |
| | **Accuracy** | | | 0.9183 |
| | **Micro-average** | 0.9183 | 0.9183 | 0.9183 |
| | | **Precision** | **Recall** | **F1-score** |
| **4th fold** | **ARGs** | 0.9975 | 0.9950 | 0.9962 |
| | **VFs** | 0.9071 | 0.8300 | 0.8668 |
| | **NSs** | 0.8414 | 0.9150 | 0.8766 |
| | **Accuracy** | | | 0.9133 |
| | **Micro-average** | 0.9133 | 0.9133 | 0.9133 |
| | | **Precision** | **Recall** | **F1-score** |
| **5th fold** | **ARGs** | 1.0000 | 0.9850 | 0.9924 |
| | **VFs** | 0.8801 | 0.8625 | 0.8712 |
| | **NSs** | 0.8623 | 0.8925 | 0.8771 |
| | **Accuracy** | | | 0.9133 |
| | **Micro-average** | 0.9133 | 0.9133 | 0.9133 |

**Table S3.** The detailed cross-validation results for each part in HyperVR-ARGs under 5-fold cross validation

| Model | Feature | Acc. | Sen. | Spec. | Prec. | MCC | AUC |
|---|---|---|---|---|---|---|---|
| Extra Trees | PSSM-Composition | 0.9828±0.0044 | 0.9490±0.0136 | 0.9997±0.0005 | 0.9994±0.0011 | 0.9616±0.0098 | 0.9985±0.0008 |
| | AADP-PSSM | 0.9906±0.0023 | 0.9735±0.0067 | 0.9992±0.0011 | 0.9984±0.0022 | 0.9790±0.0051 | 0.9973±0.0006 |
| | RPM-PSSM | 0.9796±0.0061 | 0.9395±0.0190 | 0.9997±0.0005 | 0.9994±0.0011 | 0.9545±0.0136 | 0.9982±0.0010 |
| DNN | Bit-score | 0.9913±0.0020 | 0.9815±0.0092 | 0.9962±0.0029 | 0.9924±0.0057 | 0.9805±0.0045 | 0.9949±0.0026 |
| Hybrid | All | 0.9958±0.0010 | 0.9905±0.0054 | 0.9985±0.0014 | 0.9970±0.0027 | 0.9906±0.0023 | 0.9984±0.0001 |

**Table S4.** The detailed cross-validation results for each part in HyperVR-VFs under 5-fold cross validation

| Model | Feature | Acc. | Sen. | Spec. | Prec. | MCC | AUC |
|---|---|---|---|---|---|---|---|
| RandomForest | AAC | 0.8594±0.0067 | 0.6640±0.0097 | 0.9572±0.0073 | 0.8861±0.0176 | 0.6765±0.0166 | 0.9277±0.0064 |
| | DPC | 0.8470±0.0060 | 0.5855±0.0117 | 0.9777±0.0037 | 0.9293±0.0118 | 0.6518±0.0152 | 0.9300±0.0037 |
| | DDE | 0.8483±0.0062 | 0.6195±0.0244 | 0.9627±0.0043 | 0.8928±0.0083 | 0.6509±0.0144 | 0.9203±0.0029 |
| | PAAC | 0.8491±0.0021 | 0.6405±0.0174 | 0.9535±0.0057 | 0.8735±0.0106 | 0.6516±0.0046 | 0.9205±0.0033 |
| | QSO | 0.8524±0.0063 | 0.6310±0.0180 | 0.9632±0.0033 | 0.8956±0.0089 | 0.6608±0.0151 | 0.9274±0.0032 |
| | PSSM-Com | 0.8948±0.0080 | 0.7670±0.0264 | 0.9587±0.0077 | 0.9032±0.0154 | 0.7595±0.0187 | 0.9539±0.0043 |
| | AADP-PSSM | 0.8971±0.0085 | 0.7820±0.0246 | 0.9547±0.0059 | 0.8963±0.0122 | 0.7648±0.0199 | 0.9604±0.0041 |
| | RPM-PSSM | 0.8830±0.0036 | 0.7200±0.0142 | 0.9645±0.0075 | 0.9106±0.0159 | 0.7324±0.0088 | 0.9512±0.0049 |
| Xgboost | AAC | 0.8656±0.0094 | 0.7410±0.0158 | 0.9279±0.0109 | 0.8376±0.0205 | 0.6916±0.0214 | 0.9254±0.0068 |
| | DPC | 0.8700±0.0092 | 0.7330±0.0269 | 0.9385±0.0042 | 0.8562±0.0093 | 0.7009±0.0220 | 0.9337±0.0049 |
| | DDE | 0.8718±0.0121 | 0.7330±0.0258 | 0.9412±0.0097 | 0.8619±0.0209 | 0.7053±0.0285 | 0.9335±0.0062 |
| | PAAC | 0.8656±0.0084 | 0.7400±0.0212 | 0.9284±0.0059 | 0.8381±0.0122 | 0.6915±0.0200 | 0.9267±0.0069 |
| | QSO | 0.8683±0.0095 | 0.7345±0.0223 | 0.9352±0.0124 | 0.8507±0.0234 | 0.6975±0.0221 | 0.9331±0.0043 |
| | PSSM-Com | 0.9098±0.0078 | 0.8315±0.0216 | 0.9490±0.0057 | 0.8908±0.0109 | 0.7947±0.0182 | 0.9624±0.0043 |
| | AADP-PSSM | 0.9103±0.0077 | 0.8355±0.0170 | 0.9477±0.0100 | 0.8892±0.0187 | 0.7962±0.0177 | 0.9650±0.0055 |
| | RPM-PSSM | 0.9128±0.0064 | 0.8335±0.0221 | 0.9525±0.0036 | 0.8977±0.0061 | 0.8015±0.0152 | 0.9606±0.0030 |
| ExtraTress | AAC | 0.8698±0.0055 | 0.6745±0.0136 | 0.9675±0.0062 | 0.9123±0.0151 | 0.7023±0.0135 | 0.9398±0.0055 |
| | DPC | 0.8495±0.0038 | 0.5795±0.0109 | 0.9845±0.0038 | 0.9493±0.0116 | 0.6604±0.0094 | 0.9337±0.0031 |
| | DDE | 0.8486±0.0044 | 0.5875±0.0131 | 0.9792±0.0047 | 0.9342±0.0135 | 0.6564±0.0110 | 0.9288±0.0016 |
| | PAAC | 0.8508±0.0035 | 0.6255±0.0213 | 0.9635±0.0062 | 0.8959±0.0125 | 0.6571±0.0077 | 0.9271±0.0021 |
| | QSO | 0.8635±0.0051 | 0.6435±0.0185 | 0.9735±0.0046 | 0.9241±0.0113 | 0.6889±0.0118 | 0.9400±0.0031 |
| | PSSM-Com | 0.9013±0.0087 | 0.7740±0.0287 | 0.9650±0.0080 | 0.9174±0.0159 | 0.7749±0.0200 | 0.9608±0.0042 |
| | AADP-PSSM | 0.9040±0.0085 | 0.7990±0.0228 | 0.9565±0.0049 | 0.9018±0.0109 | 0.7807±0.0199 | 0.9651±0.0042 |
| | RPM-PSSM | 0.8844±0.0082 | 0.7110±0.0224 | 0.9712±0.0086 | 0.9256±0.0207 | 0.7369±0.0199 | 0.9580±0.0043 |
| | AAC | 0.8443±0.0097 | 0.6955±0.0168 | 0.9187±0.0137 | 0.8112±0.0246 | 0.6410±0.0218 | 0.9051±0.0080 |
| | DPC | 0.8616±0.0084 | 0.7200±0.0259 | 0.9325±0.0066 | 0.8422±0.0122 | 0.6814±0.0201 | 0.9260±0.0031 |
| | DDE | 0.8585±0.0096 | 0.7135±0.0228 | 0.9309±0.0073 | 0.8380±0.0154 | 0.6739±0.0228 | 0.9242±0.0052 |
| | PAAC | 0.8500±0.0047 | 0.6945±0.0054 | 0.9277±0.0078 | 0.8280±0.0149 | 0.6536±0.0107 | 0.9121±0.0054 |

| | | | | | | | |
|---|---|---|---|---|---|---|---|
| **GradientBoosting** | **QSO** | 0.8551±0.0064 | 0.7120±0.0102 | 0.9267±0.0051 | 0.8293±0.0116 | 0.6662±0.0149 | 0.9182±0.0029 |
| | **PSSM-Com** | 0.8981±0.0047 | 0.8120±0.0205 | 0.9412±0.0070 | 0.8738±0.0113 | 0.7680±0.0114 | 0.9563±0.0035 |
| | **AADP-PSSM** | 0.8983±0.0070 | 0.8095±0.0161 | 0.9427±0.0098 | 0.8764±0.0178 | 0.7682±0.0160 | 0.9594±0.0059 |
| | **RPM-PSSM** | 0.9001±0.0065 | 0.8210±0.0254 | 0.9397±0.0079 | 0.8723±0.0128 | 0.7730±0.0156 | 0.9554±0.0035 |
| **Adaboost** | **AAC** | 0.7796±0.0098 | 0.5875±0.0349 | 0.8757±0.0175 | 0.7036±0.0220 | 0.4874±0.0238 | 0.8220±0.0136 |
| | **DPC** | 0.8331±0.0156 | 0.7080±0.0176 | 0.8957±0.0145 | 0.7728±0.0291 | 0.6180±0.0351 | 0.8902±0.0113 |
| | **DDE** | 0.8371±0.0086 | 0.7205±0.0119 | 0.8955±0.0096 | 0.7753±0.0173 | 0.6280±0.0192 | 0.8951±0.0067 |
| | **PAAC** | 0.7973±0.0059 | 0.6290±0.0219 | 0.8815±0.0079 | 0.7264±0.0108 | 0.5311±0.0152 | 0.8479±0.0069 |
| | **QSO** | 0.8036±0.0050 | 0.6520±0.0173 | 0.8795±0.0125 | 0.7306±0.0168 | 0.5482±0.0105 | 0.8500±0.0087 |
| | **PSSM-Com** | 0.8824±0.0058 | 0.8030±0.0137 | 0.9222±0.0098 | 0.8380±0.0156 | 0.7333±0.0126 | 0.9368±0.0052 |
| | **AADP-PSSM** | 0.8866±0.0067 | 0.8045±0.0172 | 0.9277±0.0141 | 0.8485±0.0241 | 0.7426±0.0146 | 0.9411±0.0052 |
| | **RPM-PSSM** | 0.8820±0.0078 | 0.7995±0.0193 | 0.9232±0.0035 | 0.8388±0.0083 | 0.7318±0.0186 | 0.9370±0.0028 |
| **DNN** | **One-Hot** | 0.8463±0.0081 | 0.6610±0.0395 | 0.9390±0.0107 | 0.8450±0.0172 | 0.6443±0.0196 | 0.9072±0.0047 |
| **Hybrid** | **All** | 0.9183±0.0044 | 0.8505±0.0208 | 0.9522±0.0069 | 0.8993±0.0115 | 0.8145±0.0103 | 0.9722±0.0040 |

**Table S5.** The genes predicted by HyperVR with >95% score in Mycobacterium tuberculosis strains

| | Rank | Gene name | Prediction score | Annotation | Evidence |
|---|---|---|---|---|---|
| **ARGs** | 1 | P9WFF9 | 1.000 | Antibiotic resistance | Uniprot |
| | 2 | P9WKD3 | 0.998 | Antibiotic resistance | Uniprot |
| | 3 | O07806 | 0.984 | Antibiotic resistance | Uniprot |
| | 4 | P9WJX5 | 0.950 | Transmembrane transport | GO-ECO:0000256 |
| | **Rank** | **Gene name** | **Prediction score** | **Annotation** | **Evidence** |
| | 1 | Q79FW5 | 1.000 | Virulence | Uniprot |
| | 2 | P9WFU7 | 1.000 | Virulence | Uniprot |
| | 3 | Q79FU2 | 1.000 | Virulence | Uniprot |
| | 4 | Q79FU0 | 1.000 | Virulence | Uniprot |
| | 5 | P0DOA7 | 1.000 | Virulence | Uniprot |
| | 6 | P0DOA6 | 1.000 | Secreted | Uniprot |
| | 7 | P9WNJ5 | 1.000 | Virulence | Uniprot |
| | 8 | P9WP43 | 1.000 | Virulence | Uniprot |
| | 9 | P9WI39 | 1.000 | Virulence | Uniprot |
| | 10 | P9WHU1 | 1.000 | Virulence | Uniprot |
| | 11 | P9WGU7 | 1.000 | Virulence | Uniprot |
| | 12 | P9WNI7 | 0.999 | Virulence | Uniprot |
| | 13 | P9WNJ3 | 0.999 | Virulence | VFDB-VFG002400(gb \| YP_177838)-100.0 |
| | 14 | I6Y9J2 | 0.998 | Peptidoglycan synthesis | Uniprot |
| | 15 | Q79G04 | 0.998 | Virulence | Uniprot |

| | | | | |
|---|---|---|---|---|
| 16 | P9WP41 | 0.998 | Virulence | Uniprot |
| 17 | P9WI47 | 0.998 | Virulence | Uniprot |
| 18 | I6Y2J4 | 0.997 | Virulence | Uniprot |
| 19 | O05442 | 0.997 | Virulence | Uniprot |
| 20 | P9WI69 | 0.997 | Virulence | Uniprot |
| 21 | L0T5T4 | 0.996 | Virulence | Uniprot |
| 22 | O53780 | 0.994 | Virulence | Uniprot |
| 23 | Q50703 | 0.994 | Virulence | Uniprot |
| 24 | Q79FB3 | 0.993 | Virulence | Uniprot |
| 25 | Q79FR5 | 0.992 | Virulence | Uniprot |
| 26 | Q79FR3 | 0.992 | Virulence | Uniprot |
| 27 | I6X486 | 0.992 | Virulence | Uniprot |
| 28 | P9WIF1 | 0.991 | Virulence | VFDB-VFG024676(gi:406030855)-62.0 |
| 29 | P9WJ67 | 0.991 | Virulence | Uniprot |
| 30 | P9WI83 | 0.990 | Virulence | Uniprot |
| 31 | P9WKQ1 | 0.989 | Cytolysis | Uniprot |
| 32 | P9WMZ9 | 0.989 | Virulence | Uniprot |
| 33 | P9WIR3 | 0.989 | Virulence | Uniprot |
| 34 | I6Y9F7 | 0.988 | Virulence | Uniprot |
| 35 | P9WI81 | 0.988 | Virulence | Uniprot |
| 36 | Q79FD3 | 0.986 | Proteolysis | GO-ECO:0000256 |
| 37 | O07747 | 0.986 | Virulence | Uniprot |
| 38 | P9WIP7 | 0.986 | Virulence | Uniprot |
| 39 | Q6MX50 | 0.985 | Virulence | Uniprot |
| 40 | P9WHZ5 | 0.985 | Virulence | PATRIC-fig\|83332.12.peg.391-68.0 |
| 41 | P96855 | 0.985 | Virulence | Uniprot |
| 42 | L7N680 | 0.984 | Virulence | Uniprot |
| 43 | Q79FI9 | 0.984 | Virulence | Uniprot |
| 44 | P9WI11 | 0.983 | Virulence | PATRIC-fig\|83332.12.peg.3735-62.6 |
| 45 | O53740 | 0.981 | Extracellular region | GO-ECO:0007005 |
| 46 | P9WFD7 | 0.981 | Virulence | Uniprot |
| 47 | I6X9F4 | 0.976 | Membrane | GO-ECO:0000323 |
| 48 | P71744 | 0.976 | Sulfate transmembrane transport | GO-ECO:0000256 |
| 49 | P9WK47 | 0.976 | Lipid transport | GO-ECO:0000323 |
| 50 | Q79FS8 | 0.975 | Virulence | Uniprot |
| 51 | Q79FI8 | 0.975 | Virulence | PATRIC-fig\|83332.12.peg.3735-68.3 |
| 52 | P9WIF5 | 0.974 | Virulence | Uniprot |
| 53 | O53651 | 0.974 | Membrane | GO-ECO:0000323 |
| 54 | P9WK37 | 0.974 | Secreted | Uniprot |

| | | | | | |
|---|---|---|---|---|---|
| | 55 | Q6MX48 | 0.973 | Virulence | PATRIC-fig∣83332.12.peg.391-62.3 |
| | 56 | P9WJ83 | 0.973 | Transmembrane transport | GO-ECO:0000364 |
| | 57 | O07750 | 0.973 | Membrane | GO-ECO:0000256 |
| | 58 | O53945 | 0.973 | Virulence | VFDB-VFG002403(gb∣NP_216312)-100.0 |
| | 59 | Q79FU3 | 0.972 | Virulence | PATRIC-fig∣83332.12.peg.2031-54.0 |
| | 60 | O33346 | 0.972 | Penicillin binding | GO-ECO:0000318 |
| | 61 | P9WHP5 | 0.972 | Virulence | Uniprot |
| | 62 | P9WMZ3 | 0.972 | Virulence | Uniprot |
| | 63 | L7N661 | 0.971 | Virulence | PATRIC-fig∣83332.12.peg.184-100.0 |
| | 64 | P9WGT9 | 0.971 | Virulence | Uniprot |
| | 65 | O53505 | 0.971 | Virulence | PATRIC-fig∣83332.12.peg.2422-100.0 |
| | 66 | I6X5W6 | 0.970 | Possible conserved membrane or exported protein | Uniprot |
| | 67 | P9WI21 | 0.970 | Virulence | PATRIC-fig∣fig83332.12.peg.391-69.6 |
| | 68 | L7N675 | 0.970 | Virulence | Uniprot |
| | 69 | P9WL77 | 0.970 | Peptide transport | GO-ECO:0000318 |
| | 70 | I6Y3Q0 | 0.970 | Virulence | Uniprot |
| | 71 | O53971 | 0.969 | Virulence | VFDB-VFG010161(gi:15609107)-100.0 |
| | 72 | P9WMY5 | 0.969 | Virulence | Uniprot |
| | 73 | I6YGJ4 | 0.968 | Kinase activity | GO-ECO:0000323 |
| **VFs** | 74 | O53489 | 0.967 | Plasma membrane | GO-ECO:0007005 |
| | 75 | P9WIS7 | 0.967 | Virulence | Uniprot |
| | 76 | P9WIF7 | 0.966 | Virulence | PATRIC-fig∣83332.12.peg.2031-67.2 |
| | 77 | Q79FP0 | 0.966 | Virulence | Uniprot |
| | 78 | O05448 | 0.966 | Cellular anatomical entity | GO-ECO:0007669 |
| | 79 | I6Y461 | 0.966 | Virulence | VFDB-VFG010046(gi:15607733)-100.0 |
| | 80 | I6XHM5 | 0.966 | Virulence | VFDB-VFG043447(gi:57116905)-56.7 |
| | 81 | P9WMB5 | 0.966 | Virulence | Uniprot |
| | 82 | P9WGT7 | 0.966 | Virulence | PATRIC-fig∣83332.111.peg.1033-64.8 |
| | 83 | P9WLL1 | 0.966 | Membrane | GO-ECO:0000323 |
| | 84 | P9WM15 | 0.966 | Extracellular region | GO-ECO:0007005 |
| | 85 | Q79G13 | 0.965 | Hydrolase activity | GO-ECO:0000323 |
| | 86 | I6YC95 | 0.964 | Virulence | VFDB-VFG010277(gi:15610630)-100.0 |
| | 87 | Q6MX04 | 0.964 | Virulence | PATRIC-fig∣83332.12.peg.3735-61.9 |
| | 88 | O50393 | 0.964 | Virulence | Uniprot |
| | 89 | I6YCA3 | 0.964 | Virulence | Uniprot |
| | 90 | P9WJC1 | 0.964 | Virulence | VFDB-VFG002389(gb∣NP_218396)-100.0 |
| | 91 | P96265 | 0.963 | Membrane | GO-ECO:0000323 |
| | 92 | I6YAY5 | 0.963 | Uncharacterized | |
| | 93 | O05582 | 0.963 | Plasma membrane | GO-ECO:0007005 |

| | | | | |
|---|---|---|---|---|
| 94 | P9WI03 | 0.963 | Virulence | VFDB-VFG029034(gi:499076672)-60.0 |
| 95 | I6X7F2 | 0.962 | Membrane | GO-ECO:0000323 |
| 96 | Q6MWW0 | 0.962 | Virulence | PATRIC-fig|83332.12.peg.391-64.4 |
| 97 | P9WHU5 | 0.962 | Virulence | PATRIC-fig|216594.6.peg.2437-83.4 |
| 98 | P9WI37 | 0.962 | Virulence | PATRIC-fig|83332.12.peg.3735-62.9 |
| 99 | P9WHY1 | 0.962 | Induction by symbiont of host immune response | GO-ECO:0000314 |
| 100 | P9WK73 | 0.962 | Membrane | GO-ECO:0000323 |
| 101 | O06295 | 0.961 | Lipoprotein | Uniprot |
| 102 | P9WK43 | 0.961 | Membrane | GO-ECO:0000323 |
| 103 | I6XEF1 | 0.960 | Virulence | PATRIC-fig|83332.12.peg.2681-60.2 |
| 104 | Q79FA8 | 0.960 | Virulence | VFDB-VFG023519(gi:333990534)-67.8 |
| 105 | P9WK45 | 0.960 | Virulence | Uniprot |
| 106 | P9WI29 | 0.960 | Virulence | PATRIC-fig|83332.12.peg.391-60.5 |
| 107 | Q79FH3 | 0.960 | Virulence | PATRIC-fig|83332.12.peg.2370-100.0 |
| 108 | Q79FY7 | 0.959 | Virulence | Victors-gi|57116756-100.0 |
| 109 | Q79FV3 | 0.959 | Virulence | PATRIC-fig|83332.12.peg.2031-65.9 |
| 110 | L0TCB8 | 0.959 | Virulence | PATRIC-fig|83332.12.peg.2031-59.8 |
| 111 | Q10690 | 0.959 | Induction by symbiont of host immune response | GO-ECO:0000314 |
| 112 | P9WK65 | 0.959 | Virulence | Uniprot |
| 113 | P71986 | 0.958 | Extracellular region | GO-ECO:0007005 |
| 114 | O06155 | 0.957 | Beta-lactamase activity | GO-ECO:0000256 |
| 115 | O05445 | 0.957 | Plasma membrane | GO-ECO:0007005 |
| 116 | L7N693 | 0.957 | Virulence | PATRIC-fig|83332.12.peg.2031-62.2 |
| 117 | O05444 | 0.957 | Membrane | GO-ECO:0000323 |
| 118 | P9WP39 | 0.957 | Hydrolase activity | GO-ECO:0000323 |
| 119 | Q50702 | 0.957 | Plasma membrane | GO-ECO:0007005 |
| 120 | P9WGU1 | 0.957 | Virulence | Uniprot |
| 121 | I6YGB1 | 0.956 | Virulence | Victors-gi|15610633-100.0 |
| 122 | P95206 | 0.956 | Extracellular region | GO-ECO:0007005 |
| 123 | P9WJ71 | 0.956 | Extracellular region | GO-ECO:0007005 |
| 124 | P9WK69 | 0.956 | Membrane | GO-ECO:0000323 |
| 125 | O07726 | 0.955 | Extracellular region | GO-ECO:0007005 |
| 126 | O69622 | 0.955 | Virulence | Uniprot |
| 127 | P9WJA1 | 0.955 | Interaction with host | GO-ECO:0000315 |
| 128 | Q79FP3 | 0.954 | Virulence | PATRIC-fig|83332.12.peg.2031-63.0 |
| 129 | L7N667 | 0.954 | Virulence | VFDB-VFG023529(gi:183982680)-62.2 |
| 130 | I6Y1I5 | 0.954 | Membrane | GO-ECO:0000323 |
| 131 | O53968 | 0.954 | Virulence | VFDB-VFG010093(gi:15609104)-100.0 |
| 132 | O07745 | 0.954 | Extracellular region | GO-ECO:0007005 |

| | | | | |
|---|---|---|---|---|
| 133 | I6YGS7 | 0.954 | Virulence | VFDB-VFG024677(gi:379754765)-70.7 |
| 134 | P9WNU9 | 0.954 | Virulence | Uniprot |
| 135 | L7N697 | 0.953 | Virulence | PATRIC-fig\|83332.12.peg.2681-60.0 |
| 136 | O05854 | 0.953 | Plasma membrane | GO-ECO:0007005 |
| 137 | O53963 | 0.953 | Lipoprotein | Uniprot |
| 138 | O53638 | 0.953 | Peptidoglycan biosynthetic process | GO-ECO:0000322 |
| 139 | P9WI15 | 0.953 | Virulence | PATRIC-fig\|83332.12.peg.3735-52.6 |
| 140 | P9WLR9 | 0.953 | Peptidoglycan-based cell wall | GO-ECO:0007005 |
| 141 | I6Y7L4 | 0.952 | Virulence | PATRIC-fig\|83332.12.peg.391-100.0 |
| 142 | O53206 | 0.951 | Uncharacterized | |
| 143 | O53200 | 0.951 | Membrane | GO-ECO:0000323 |
| 144 | P9WKS1 | 0.951 | Plasma membrane | GO-ECO:0007005 |
| 145 | Q79FL8 | 0.950 | Virulence | Uniprot |
| 146 | O05859 | 0.950 | Zinc ion binding | GO-ECO:0000256 |
| 147 | L7N659 | 0.950 | Virulence | VFDB-VFG024676(gi:406030855)-60.0 |
| 148 | P9WK67 | 0.950 | Membrane | GO-ECO:0000323 |

### SUPPLEMENTARY FIGURES

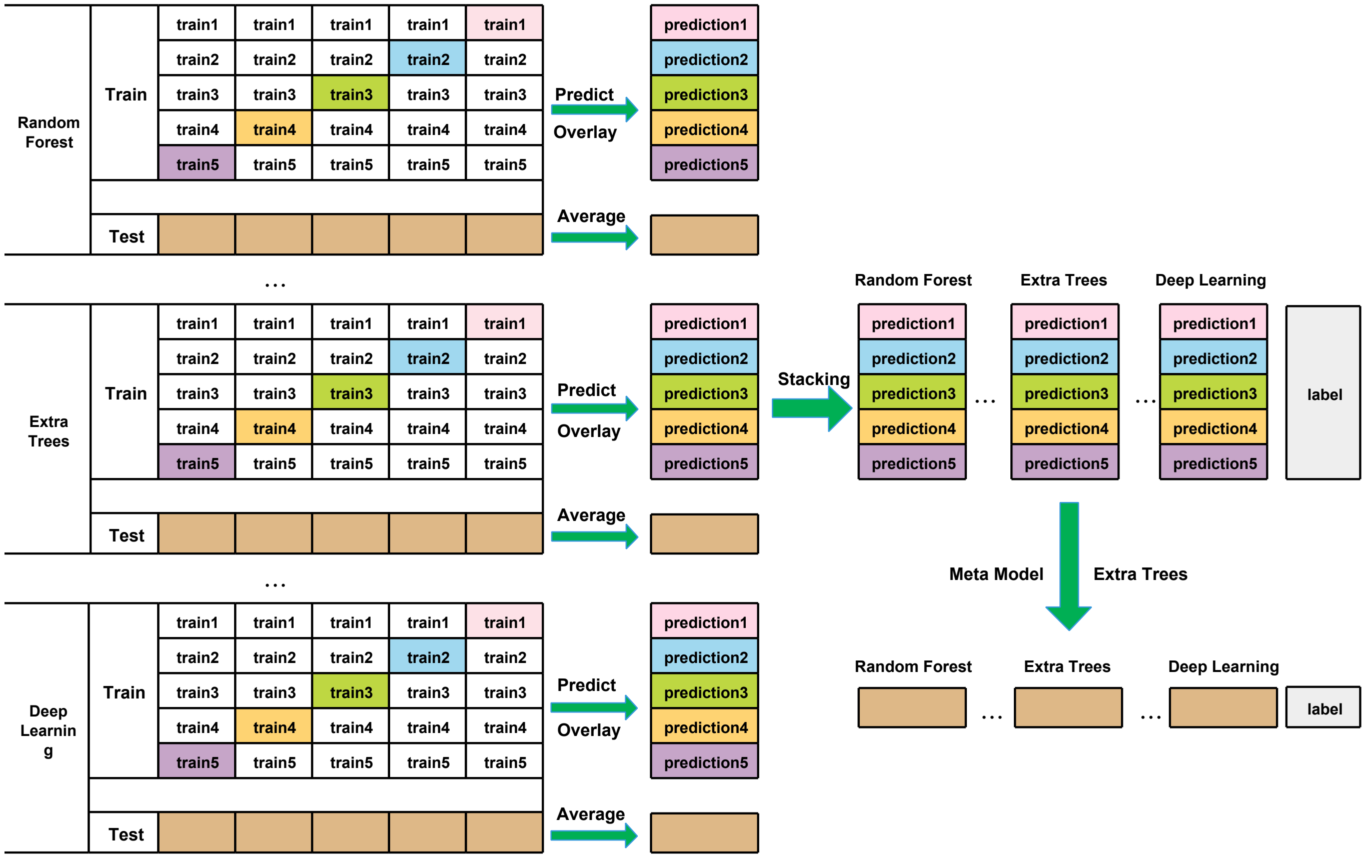


**Fig. S1.** The detailed process for HyperVR using a stacking strategy

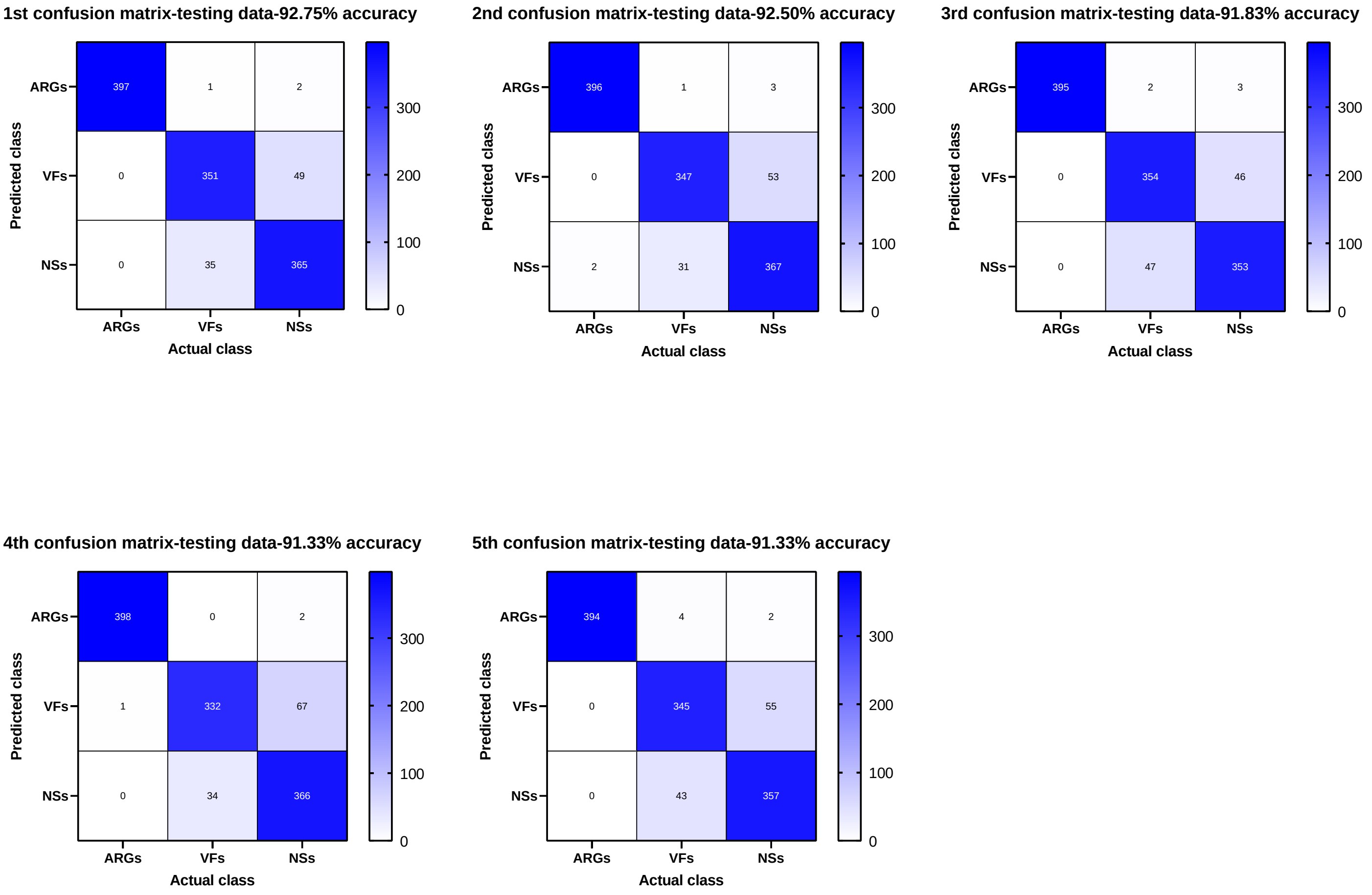


**Fig. S2.** The detailed confusion matrices of HyperVR under 5-fold cross-validation

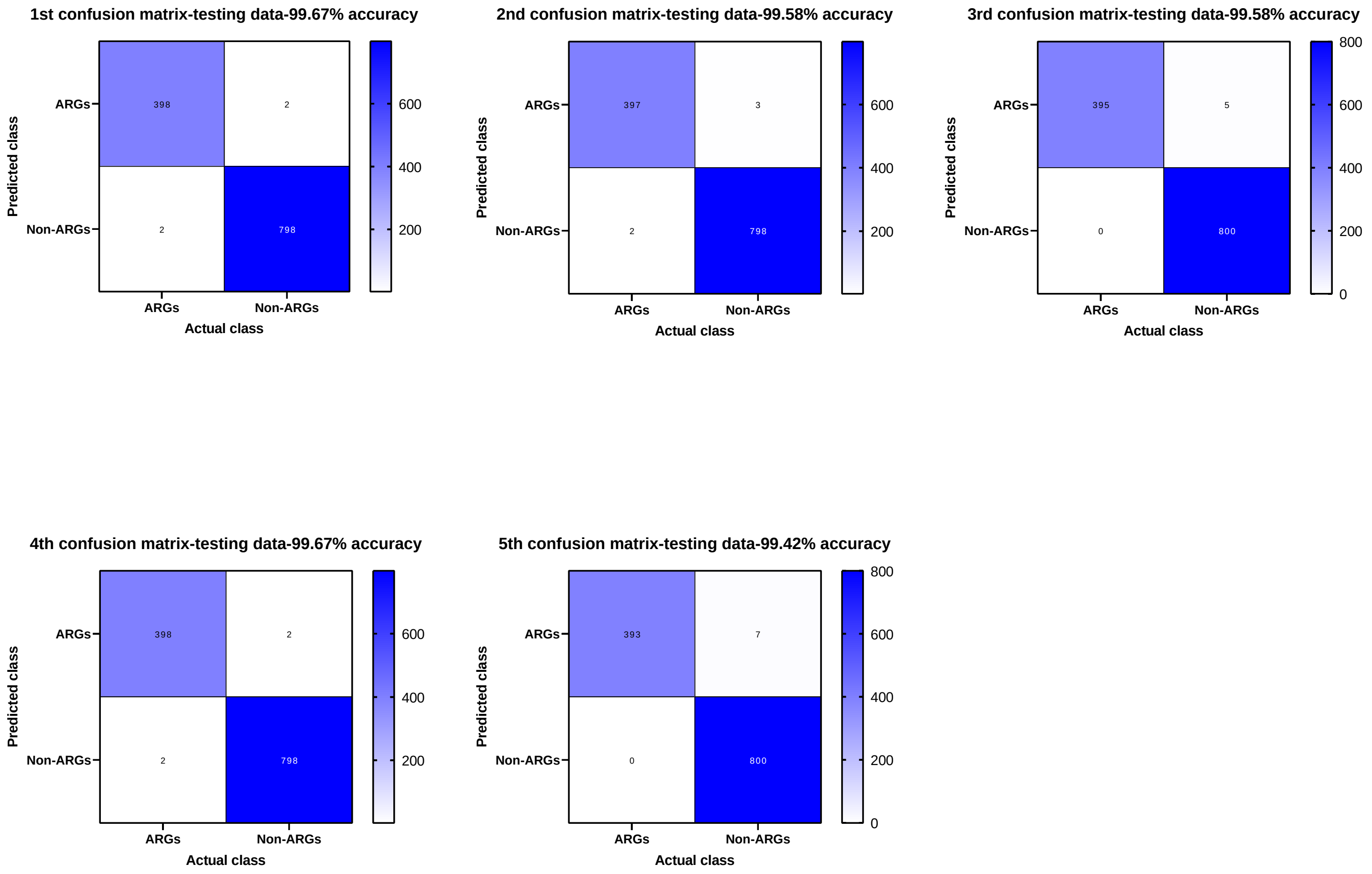


**Fig. S3.** The detailed confusion matrices of HyperVR-ARGs under 5-fold cross-validation

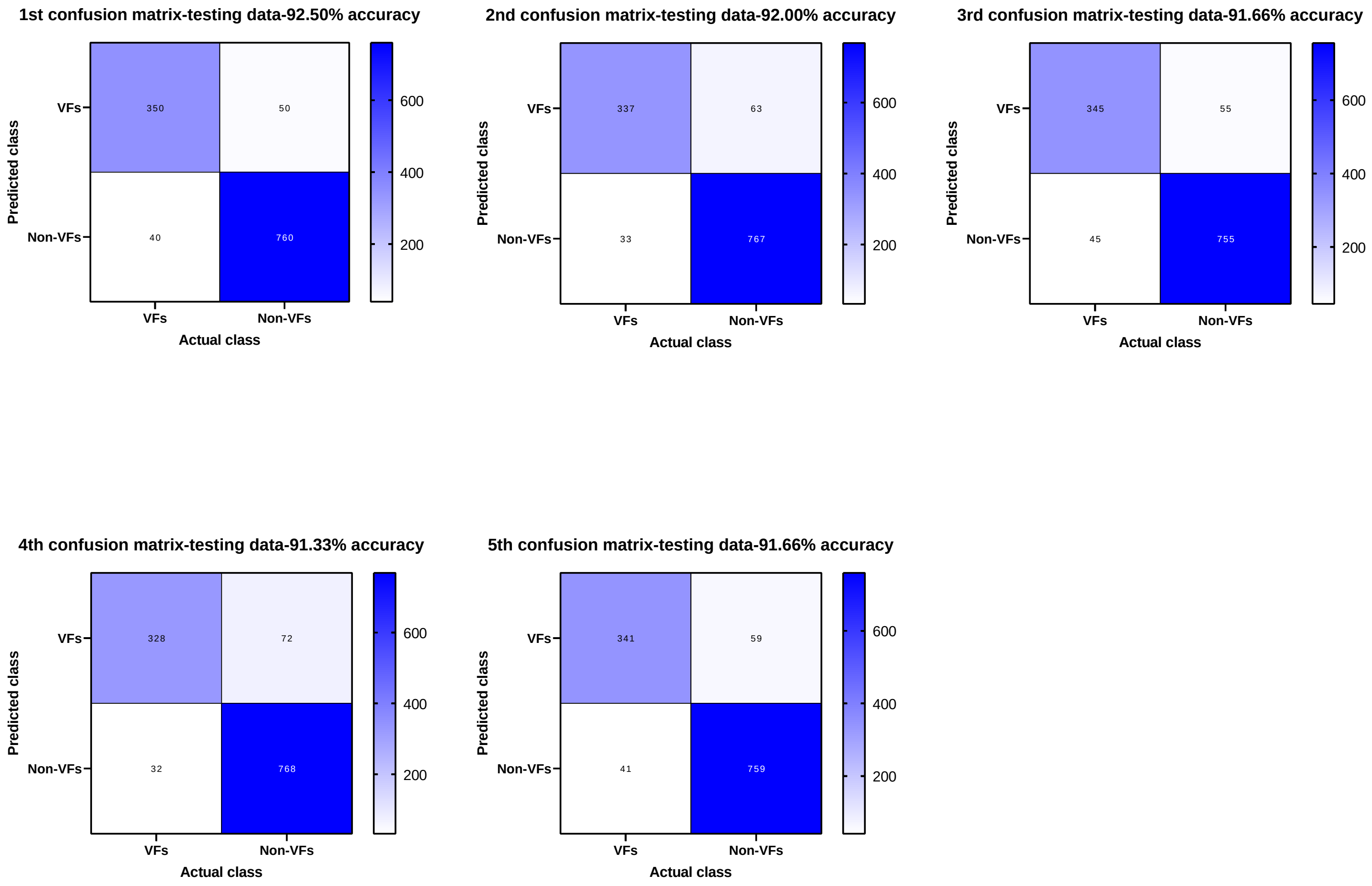


**Fig. S4.** The detailed confusion matrices of HyperVR-VFs under 5-fold cross-validation
